# A Novel Tiled-Amplicon Sequencing Assay Targeting the *Tomato Brown Rugose Fruit Virus* (*ToBRFV*) Genome Reveals Widespread Distribution in Municipal Wastewater Treatment Systems in the Province of Ontario, Canada

**DOI:** 10.1101/2023.11.02.565359

**Authors:** Delaney Nash, Isaac Ellmen, Jennifer J. Knapp, Ria Menon, Alyssa K. Overton, Jiujun Cheng, Michael D.J. Lynch, Jozef I. Nissimov, Trevor C. Charles

## Abstract

*Tomato Brown Rugose Fruit Virus* (*ToBRFV*) is a plant pathogen that infects important *Solanaceae* crop species and can dramatically reduce tomato crop yields. The *ToBRFV* has rapidly spread around the globe due to its ability to escape detection by antiviral host genes, most notably *Tm-2^2^*, which are used to confer resistance to other *Tobamoviruses* in tomato plants. Development of robust and reproducible methods for detecting viruses in the environment aids in the tracking and reduction of pathogen transmission. We detected *ToBRFV* in municipal wastewater influent (WWI) samples, likely due to its presence in human waste, demonstrating a widespread distribution of *ToBRFV* in WWI throughout Ontario, Canada. To aid in global *ToBRFV* surveillance efforts, we developed a tiled-amplicon approach to sequence and track the evolution of *ToBRFV* genomes in municipal WWI. Our assay recovers 97.5% of the 6393 bp *ToBRFV* RefSeq genome, omitting the terminal 5’ and 3’ ends. We demonstrate that our sequencing assay is a robust, sensitive, and highly specific method for recovering *ToBRFV* genomes. Our *ToBRFV* assay was developed using existing ARTIC Network resources, which includes genome specific primer design, sequencing library prep, and read analysis. Additionally, we adapted our lineage abundance estimation tool, Alcov, to estimate the abundance of *ToBRFV* clades in samples.

## Introduction

### Tomato Brown Rugose Fruit Virus global incidence, impact, and phylogeny

In Canada, fresh market tomatoes have been the most abundantly produced greenhouse vegetable since record keeping began in 1955. In 2022, >293M kg of fruit with a farm gate value of >$793M were produced, with >213M kg of fruit with a farm gate value of >$540M produced in Ontario (*Statistics Canada*, n.d.). The *Tomato Brown Rugose Fruit Virus* (*ToBRFV*) is a devastating plant-pathogen that infects tomato and pepper plants and can reach nearly 100% disease incidence causing massive crop losses and placing a high economic burden on farmers (Salem et al., 2016; Jones, 2021; Zhang, et al. 2022). Thus, the control or elimination of this virus is a major obstacle in maintaining secure global tomato crop yields.

The first incidence of *ToBRFV* was reported by Salem and colleagues (2016) following a 2015 outbreak in Jordan. Shortly thereafter, Luria and colleagues (2017) identified *ToBRFV* as the causative agent of a 2014 tomato disease outbreak in Israel. It has spread to at least 35 countries since, including major tomato producing countries such as China, Türkiye, and Mexico (Faostat, 2020; Fidan et al., 2021; Ma et al. 2021; Zhang et al. 2022). Global transmission and clade specific mutations of *ToBRFV* strains are monitored through a curated NextStrain database updated by authorized curators with each updated version referred to as a build (Botermans et al., 2023; van de Vossenberg, et al. 2020).

### Taxonomic classification, distinguishing between species, lineages, and strains

The *ToBRFV* is a member of the *Tobamovirus* genus, a group of highly-virulent plant pathogens that includes 37 virus species with diverse and distinct host-ranges (Jones, 2021; ICTV, 2017; Gibbs, 1999). ICTV species demarcation criteria for said genus is defined as <90% shared nucleotide identity between related genomes, while strains of the same species share >90% nucleotide sequence identity (Adams et al., 2009; Lefkowitz, et al. 2018; Abrahamian et al., 2022). The *ToBRFV* genome shares the highest sequence identity with *Tobacco Mosaic Virus (TMV)* isolate Ohio V at 82.4% shared identity, and fulfills species demarcation criteria (Salem et al., 2016; Ochar et al., 2023). All *ToBRFV* isolates available in NextStrain share greater than 90% sequence identity, meeting the demarcation criteria (data not shown). The *ToBRFV* 2022 NextStrain build (version 3) contains 179 *ToBRFV* isolates grouped into eight distinct clades which represent diverging lineages of *ToBRFV* strains (Botermans et al., 2023; van de Vossenberg, et al. 2021).

### ToBRFV gene content, mode of infection, and host immune-escape

*Tobamovirus* virions are rod-shaped particles up to 300 nm long, 20 nm wide, and are non-enveloped (Adams et al., 2009; Adams et al., 2017; Ochar et al., 2023). The genome structure and gene order is conserved amongst all *Tobamoviruses*, consisting of a ∼6.4 kbp monopartite single-stranded positive-sense RNA genome containing four open reading frames (ORFs) (Adams et al., 2017; Ochar et al., 2023). Replication proteins are encoded by ORF1 & ORF2, whereas virus structural proteins are encoded by ORF3 & ORF4 and expressed through subgenomic RNA (Zhang et al. 2022). ORF1 encodes a 126 kDa RNA-dependent RNA polymerase (RdRp), while ORF2 encodes a 183 kDa helicase protein through an amber UAG stop codon downstream of ORF1 (Salem et al., 2016). ORF3 encodes a 30 kDa movement protein (MP) that facilitates cell-to-cell transmission through the plant plasmodesmata. The MP binds to multiple cellular components that transport the viral RNA through the plasmodesmata. Lastly, ORF4 encodes a coat protein which encapsulates the virus genome forming a thin rod-like structure (Adams et al., 2009; Salem et al., 2016).

Changes in the *ToBRFV* MP sequence break *Tobamovirus-*resistance genes that have been introduced into most commercial crop species to provide protection against *Tobamoviruses* (Hak & Spiegelman, 2021). *ToBRFV* MP amino acid mutations result in structural changes that prevent MP binding by the plant *Tm-2^2^* receptor protein, thereby preventing activation of the hypersensitive response and leading to severe infection symptoms (Hak & Spiegelman, 2021). Recently, *ToBRFV-*resistant tomato seed-lines have been commercially developed, however, only a limited number of varieties are currently available (https://www.vegetables.bayer.com; https://www.syngentavegetables.com/resources/ToBRFV).

### The ToBRFV NextStrain Database and Phylogeny

The NextStrain database is a useful resource for scientists and policy makers to understand transmission routes and establish disease control measures (van de Vossenberg et al., 2020). For example, genomic sequence data from commercial seeds was analyzed using NextStrain phylogenies which alerted Dutch authorities to the illegal use of a *ToBRFV* cross-protective species (Botermans et al., 2023). *ToBRFV* genome sequences can be submitted and incorporated into updated builds by authorized curators and this community-based approach has increased the number of genomes available in each version (Botermans et al., 2023; van de Vossenberg et al., 2021). The *ToBRFV* 2019, 2020, and 2022 NextStrain builds contain 63, 118, and 179 *ToBRFV* strains respectively which can be phylogenetically grouped into three, six, and eight clades respectively, representing diverging *ToBRFV* clades (Botermans et al., 2023; van de Vossenberg, et al. 2021; van de Vossenberg et al., 2020). However, when Abrahamian and colleagues (2022) analyzed genome sequences of 123 *ToBRFV* strains retrieved from GenBank as well as another 22 isolated from plant tissues, they identified three distinct clades. Zhang and colleagues (2022) also phylogenetically analyzed 78 *ToBRFV* genomes and identified three distinct clades. These differences between identified clades are likely a result of the differing phylogenetic analysis tools and genome sequences used by each study. Additionally, the majority of isolated strains originate from the Netherlands and this sampling bias has likely skewed *ToBRFV* phylogenetic analyses (Abrahamian et al., 2022; Botermans et al., 2023).

To fully understand *ToBRFV* phylogeny, an increased number of genome sequences from various regions must be obtained, which can be aided by improved sequencing methods (Abrahamian et al., 2022; Zhang et al., 2022). To aid in *ToBRFV* identification and surveillance efforts, we developed a robust and specific *ToBRFV* genome tiled-amplicon sequencing assay utilizing short-read Illumina sequencing technology (Figure 1).

**Figure 1.**
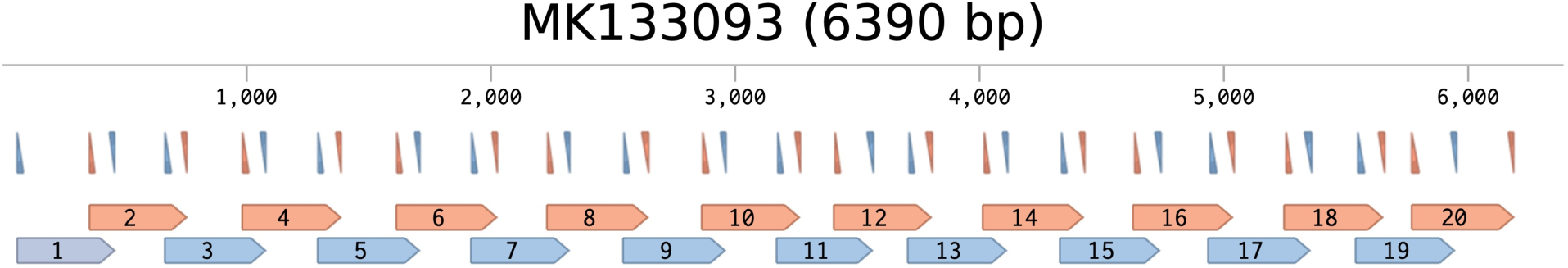
*ToBRFV*-specific primer binding sites and resulting amplicons mapped to the *ToBRFV* MK133093 genome sequence. *In silico* visualization of primer binding sites; forward and reverse primers are denoted by the upward and downward pointing triangles, respectively, each resulting amplicon is numbered and denoted by blue and orange flags (Benchling, 2023). Pool 1 primers and amplicons are blue, pool 2 primers and amplicons are orange. Nucleotide positions are labeled at 1000 bp intervals.

### Use of PCR-enrichment sequencing assays for viral detection

High-throughput shotgun sequencing is capable of producing millions of reads from a single sample (Bai et al., 2022; Pérez-Losada et al., 2020; Sandybayev et al., 2022). However, the low-abundance of viral reads and the high-abundance of host and background reads in environmental samples can make it challenging to sequence viral genomes without utilizing virus-specific enrichment methods such as PCR (Pérez-Losada et al., 2020; Yang et al., 2018). PCR-enrichment assays are often used to amplify specific DNA sequences; however, they require the development of target-specific primers (Kumar et al., 2017; Quick et al., 2017).

Recently, an open-source collaboration called the ARTIC Network has created adaptable user-friendly tools to develop virus-specific sequencing assays utilizing PCR-enrichment (https://artic.network/). The ARTIC Network was originally developed for monitoring the transmission and evolution of viruses such as MERS-CoV and Ebola but has since been adapted to streamline the design of species-specific tiled-amplicon sequencing assays (Quick et al., 2017; ARTIC Network, 2023). ARTIC primer schemes are designed to target short overlapping fragments, referred to as amplicons, that span a specific genome of interest (Figure 1). PCR-enriched amplicons can be sequenced using either Illumina or Nanopore platforms to obtain complete genome sequences. ARTIC primer schemes have been used to target many viral genomes, including but not limited to *Zika, Ebola, Dengue,* and *SARS-CoV-2* (Quick et al., 2017; Kareninen et al., 2019, Hill et al., 2018; ARTIC Network, 2020; ARTIC Network, 2023).

We utilized the ARTIC infrastructure to develop a nearly whole genome tiled-amplicon sequencing assay that enriches *ToBRFV* genomes for Illumina sequencing. During our ongoing exploration of SARS-CoV-2 content in wastewater influent (WWI), we observed widespread occurrence of the *ToBRFV* sequences in samples from multiple treatment plants in Ontario, Canada (data available upon request). Indeed, several recent studies have observed *ToBRFV* sequences in WWI, indicating this virus is likely wide-spread and prevalent in wastewater systems in many countries (Bačnik et al., 2020; Mehle et al., 2023; Rothman et al., 2022).

### *ToBRFV* Occurrence and Transmission in Wastewater

*ToBRFV* sequences have been observed in wastewater samples from Slovenia and California (Bačnik et al., 2020; Mehle et al., 2023, Rothman & Whiteson, 2022). In 2020, Bačnik and colleagues demonstrated wastewater influent contains active *ToBRFV* virions which, when concentrated, can cause asymptomatic infections in Tomato plants.

Reclaimed wastewater is commonly used to irrigate crops, however several *Tobamovirus* species, including *PMMoV*, remain transmissible in treated wastewater effluent (Bačnik et al., 2020). Thus, *ToBRFV* could potentially be transmitted to crops via reclaimed wastewater during irrigation. Moreover, in 2023, Mehle and colleagues demonstrated that *ToBRFV*-infected plants shed active virions through their roots which can then spread to other plants through shared hydroponic systems that distribute nutrient solution. This suggests that plants exposed to ToBRFV contaminated wastewater could potentially become infected and cause wide-spread outbreaks. This could be a challenge for growers who utilize reclaimed wastewater to irrigate crops (Christou et al., 2017; Goh et al., 2022). However, it has not yet been shown that *ToBRFV* virions remain infective after conventional wastewater treatment methods. Thus, the possibility remains that treated wastewater effluent is suitable for irrigation and further work must be performed to determine *ToBRFV* transmissibility (Bačnik et al., 2020; Mehle et al., 2023).

The entry mode of *ToBRFV* virions into wastewater has not yet been established. One possible explanation for *ToBRFV*-transmission is through environmental sources such as waste from regional greenhouses and fields with on-going *ToBRFV* infections (Rothman & Whiteson, 2022). Another possible mode of transmission could be through a human dietary-route (Rothman & Whiteson, 2022). If the virus is present in produce and tomato or pepper-based products, then the excretion of these products by humans could transmit *ToBRFV* virions into WWI (Zhang et al., 2006). To test these hypotheses, WWI could be obtained and tested for *ToBRFV* from regions without *ToBRFV*-infections or where there is no tomato or pepper cultivation to eliminate the possibility of transmission through agricultural waste. Additionally, tomato and pepper products from grocery stores, as well as human feces, could be tested for the *ToBRFV* to establish a dietary-route of transmission (Zhang et al., 2006). Similarly, a dietary-mode of *PMMoV* transmission to human feces has been established, thus, the human-dietary mode of *ToBRFV* transmission into WWI is the more probable scenario (Zhang et al., 2006).

### *ToBRFV* Detection and Sequencing

Current *ToBRFV* screening procedures, such as RT-qPCR and loop-isothermal amplification assays, can be used to determine the presence or absence of *ToBRFV* sequences, quantify the viral load, and provide more rapid results than sequencing assays. However, many of these assays cannot detect strain-specific mutations or they require development of specialized strain-specific primers/probes to track transmission routes and determine sources of infection (Mehle et al., 2023; Natarajan et al., 2023; Rizzo et al., 2021; Zhang et al., 2022). Moreover, increased evolutionary pressure from disease eradication efforts could result in the emergence of novel mutations which evade or reduce the effectiveness of current rapid presence/absence detection assays and eradication measures (Abrahamian et al., 2022). Thus, sequencing of complete *ToBRFV* genomes is essential to monitor virus transmission and evolution, predict emerging threats, and fully understand *ToBRFV* phylogeny (Botermans et al., 2023). However, high-throughput shotgun sequencing analysis is not ideal for recovery of *ToBRFV* genomes due to the low recovery of viral sequences, high-costs, and length of sample processing time (Zhang et al., 2022). Our tiled-amplicon assay provides a reliable cost-effective method to sequence *ToBRFV* genomes as well as aid in global surveillance efforts and uncover novel phylotypes of clades.

To aid in *ToBRFV* surveillance efforts we utilized wastewater RNA extracts from five Ontario WWI collection sites to develop and validate a novel *ToBRFV* tiled-amplicon sequencing (*ToBRFV*-Seq) assay. We prepared RNA sequencing libraries using our ARTIC-style *ToBRFV-*seq assay and an RNA shotgun sequencing assay using the NebNext Ultra II RNA Prep kit (*New England Biolabs*, E7770S). By evaluating the recovery of *ToBRFV* reads by each method we demonstrate our assay is a highly specific and efficient method for *ToBRFV* genome sequencing. Moreover, we adapted our in-house SARS-CoV-2 lineage abundance estimation tool, Alcov (Ellmen et al., 2021), to identify and estimate the relative-abundance of *ToBRFV* clades. Our adapted tool, named Altob (https://github.com/Ellmen/altob), was able to analyze clade-specific mutations and estimate the abundance of *ToBRFV* NextStrain clades 1-4, and 6-8 in samples.

## Materials and Methods

### Viral RNA Extraction from Wastewater Influent

Wastewater influent (WWI) samples were collected in plastic bottles and shipped in coolers with ice packs from five wastewater treatment plants in Ontario, Canada (Table S1). Samples from five sites (designated A-E) were processed using virion capture Nanotrap Microbiome A particles (*Ceres Nanosciences,* #44202), then RNA was extracted using the *QIAgen* RNeasy Mini kit (#74004).

Briefly, bottles of WWI were inverted to mix and allowed to settle for 1 min at room temperature, then 10 mL of wastewater supernatant containing suspended biosolids was moved to a fresh tube. A negative control sample was prepared alongside extracts using 10 mL of nuclease-free DI water instead of WWI. The supernatant was mixed with 100 µL of ER2 solution by briefly vortexing. Then, 150 µL of Microbiome A particles were added and the samples were inverted to mix. The samples were then incubated at room temperature for 10 min, with three inversions after 5 min of incubation. Samples were placed on a 15 mL tube magnetic stand; the cleared supernatant was discarded, and magnetic beads were retained.

Beads were gently resuspended in 1 mL of nuclease-free water and transferred to a sterile 1.5 mL microfuge tube. Beads were placed on a magnetic stand and incubated for 1 min, the supernatant was discarded without disturbing the bead pellet. Beads were resuspended 700 µL of RLT lysis buffer containing 1% 2-ME (*BioShop,* MER002.500) by pipetting then vortexing. Samples were incubated at room temperature for 10 min, then, 600 µL of the supernatant from each sample was manually extracting using the *QIAgen* RNeasy Mini kit (#74116). RNA was eluted in 80 µL of RNase-free water, the concentration of DNA and RNA was analyzed on a Qubit 4.0 fluorometer (*Thermofisher*, #Q33238). DNA was degraded using the Ambion DNase kit (*Thermofisher,* #AM2222), samples were incubated with 4 units of DNase at 37°C for 45 minutes. RNA was recovered using the RNeasy Mini Kit RNA clean up protocol, DNA and RNA concentration was then determined by Qubit 4.0 analysis to ensure RNA integrity and complete removal of DNA.

### Primer design and pooling

Primers were designed by first generating a multiple sequence alignment (MSA) from the genomes of 118 *ToBRFV* strains listed on the 2020 NextStrain Build (version 2), and sequences were obtained from NCBI (Table S2). A consensus sequence was generated from the MSA and uploaded to the web-based tool, PrimalScheme, to generate primer sequences targeting overlapping amplicons of approximately 400 base pairs (bp) in length. PrimalScheme generated 20 amplicons for a total of 40 primers that cover 95.7% of the *ToBRFV* genome NC_028478.1, which was used for downstream read mapping analysis. Primers were synthesized by Integrated DNA Technologies. Primers amplifying odd and even numbered amplicons were diluted and mixed in two separate aliquots (pool 1 and 2, respectively) to avoid mis-priming between overlapping amplicons (Figure 1). Each primer was pooled at a concentration of 0.5 µM in respective pools (Table S3).

### ToBRFV-Targeted Tiled-Amplicon Sequencing Library Preparation

RNA extracts from WWI samples were reverse transcribed to synthesize cDNA using the LunaScript SuperMix RT kit (*New England Biolabs,* E3010L); 10 µL of each RNA extract was used in a total reaction volume of 20 µL. First strand cDNA synthesis products were PCR-amplified in two reaction mixtures containing primer pool 1 or 2 (Figure 1). The PCR was performed using Q5 2× Master Mix (*New England Biolabs,* M0492S), 6 µL of cDNA synthesis reaction, and 2.5 µL of one primer pool, adding ddH2O to a total volume of 25 µL. Samples were initially denatured at 98°C for 30 s, followed by 35 cycles of 98°C for 15 s, 63°C for 30 s, and 72°C for 30 s, ending with a final extension of 5 min at 72°C.

PCR products were prepared for sequencing using the Illumina DNA Prep kit (#20060060) using half of the manufacturer recommended volumes. Briefly, PCR products were purified using AMPure XP beads (*Beckman Coulter,* A63881), then assessed by Qubit 4.0 fluorometer and agarose gel electrophoresis. Purified PCR products were tagmented using on-bead tagmentation and barcoded with custom unique dual indexes using six PCR cycles. Indexed PCR-products of ∼400 bp were isolated using double-sided magnetic-beads. The purified fragments were assessed using a Qubit 4.0 fluorometer and agarose gel electrophoresis.

### RNA Shotgun Sequencing Library Preparation

RNA extracts from WWI were prepared for RNA sequencing using the NEB Ultra II RNA Sequencing kit (*New England Biolabs*, E7770S), following manufacturer’s recommended protocol, section 5. RNA extracts were quantified using a Qubit 4.0 fluorometer, then, reversed-transcribed using first- and second-strand synthesis enzymes. AMPure XP beads were used to purify double-stranded cDNA fragments. Fragments were end-prepped by dA-tailing enzymes, then adaptor-ligated. Ligated fragments were purified using AMPure XP beads, PCR-enriched and barcoded with unique dual indexes (*Illumina*, 20027213) using six PCR cycles. PCR-products were purified using AMPure XP beads then assessed using a Qubit 4.0 fluorometer and agarose gel electrophoresis.

### Library Pooling and Illumina NextSeq Sequencing

*ToBRFV*-Seq and RNA shotgun prepared libraries were pooled. Samples prepared using the same method, either RNA shotgun or *ToBRFV*-Seq, were pooled at equal concentrations and shotgun libraries were pooled at 15× the concentration of *ToBRFV*-Seq libraries. The pooled libraries were sequenced on an Illumina NextSeq2000 using 2×300 sequencing and P1 chemistry. Approximately 65% of the flowcell sequencing capacity was dedicated to these samples, for a total expected yield of 52 GB.

### Read Processing, Taxonomic Classification, and ToBRFV Genome Alignment

Sequencing read quality and adaptor content was evaluated using FastQC v0.12.1 (http://www.bioinformatics.babraham.ac.uk/projects/fastqc/). Low quality reads and adaptor sequences were removed and reads were paired using Trimmomatic v0.39 using flags ILLUMINACLIP: <a>:2:30:7:2:TRUE, LEADING:3, TRAILING:3, SLIDINGWINDOW:5:15, MINLEN:125. (Bolger et al., 2014). RNA shotgun reads were trimmed using NebNext Illumina adaptor read sequence 1 AGATCGGAAGAGCACACGTCTGAACTCCAGTCA and read sequence 2 AGATCGGAAGAGCGTCGTGTAGGGAAAGAGTGT while *ToBRFV*-Seq reads were trimmed using the Illumina DNA Prep adaptor sequence CTGTCTCTTATACACATCT. The overall quality of read trimming was evaluated using FastQC v0.12.1. Read pairs passing quality control filters were taxonomically classified with Kraken2 v2.0.7-beta, using the flag -paired and the standard database, reports of read taxonomic rankings were generated for each sample (Wood et al., 2019). Quality controlled read pairs were aligned to the *ToBRFV* genome NC_028478.1 using bowtie2 v2.3.5 with the flag -local (Langmead & Salzberg, 2012). Alignments were converted into bam files with samtools v1.9 files and anvio-7.1, and then visualized using Tablet 1.21.02.08 (Danecek et al., 2021; Eren et al., 2021; Milne et al., 2013).

### Altob Implementation and Synthetic Read Simulation

Our in-house SARS-CoV-2 variant of concern lineage abundance estimation tool, Alcov, was adapted to identify and estimate *ToBRFV* clades in Illumina paired reads (Ellmen et al., 2021). We used Seaview to compute a multiple sequence alignment (MSA) with the MUSCLE algorithm using NC_028478.1 as the root genome and 124 unique genomes listed on NextStrain (Table S2) (Botermans et al., 2023; Edgar, 2004; Gouy et al., 2021). Then, a phylogenetic Maximum Likelihood tree was generated using a GTR model and 100 bootstrap replicates and compared to the 2022 NextStrain *ToBRFV* phylogenetic tree (Figure S1). Phylogenetically grouped isolates were used to generate MSAs for each clade using NC_028478.1 as the root and the MUSCLE algorithm. MSAs were used to generate a list of mutations that appear in at least 50% of clade-specific isolates, which were then used to construct constellations of clade-specific mutations to estimate *ToBRFV* clade abundances using Altob. Positions in our mutation list are relative to position one of the reference genome, whereas, the mutations reported in the NextStrain phylogeny are relative to position one of their *ToBRFV* MSA which is either 106 or 107 bp upstream of our start site.

Altob was adapted from our in-house tool Alcov by modifying the original mutation constellations and reference genome; all other functions and parameters were maintained. A detailed description of Alcov and its functions can be found in our preprint (Ellmen et al., 2021). Briefly, Altob scans sites with a minimum coverage of 40 reads for the provided mutations and calculates frequencies as the number of mutations over the total reads at that site (Ellmen et al., 2021). Altob computes the clade abundance which best explains the observed pattern of mutations utilizing linear programming, a well-studied optimization technique (Ellmen et al., 2021; Rencher & Christensen, 2012).

Synthetic read data used to benchmark Altob was generated using DGWSIM v0.1.13 and genomes of eight randomly selected *ToBRFV* strains, each belonging to a different NextStrain clade (Table S2) (Botermans et al., 2023; github.com/nh13/DWGSIM). Three datasets containing either 1 000, 10 000, or 100 000 paired 2×250 Illumina reads were generated for each of the eight genomes without introducing random mutations or random DNA reads. Files containing 100 000 paired reads were combined to generate datasets with a mixture of clade-specific mutations. An equal number of reads from eight, four, and two genomes were combined, thus, each clade is represented by 12.5%, 25%, or 50% of reads, respectively. Altob is publicly available on GitHub (https://github.com/Ellmen/altob).

## Results and Discussion

### *In silico* Primer Binding Analysis of *ToBRFV* Genome Sequences of 179 Strains

To ensure our 40 *ToBRFV*-specific primers would inclusively target a wide-variety of *ToBRFV* strains without compromising species-specific stringency, we performed an *in silico* primer binding analysis using the sequence analysis software *Benchling* (2023) and 191 *Tobamovirus* genomes. A list of 179 *ToBRFV* strain accession numbers from the NextStrain *ToBRFV* build (2022, version 3) was obtained, representing 125 unique sequences, and genomes were downloaded from NCBI. In addition, 12 *Tobamovirus* RefSeq genomes of species closely related to the *ToBRFV* were obtained from NCBI (accessed April 20th 2023) and all 191 genome sequences were imported into *Benchling* (Gibbs et al., 2018; Ochar et al., 2023). Benchling was used to probe all 191 genome sequences for potential primer binding sites using the program’s *Find Primer Binding Sites* tool. A successful match required a minimum of 18 matching bases, a maximum of 3 mismatches, and a maximum of 1 consecutive mismatched nucleotide. The positions of matching primers were mapped to each genome and visualized using *Benchling* (data not shown) to ensure the overlapping amplicon primer scheme was maintained.

All 40 *ToBRFV-*specific primers had one matching binding site within each of the 179 *ToBRFV*-strain genomes (Table 1), with between zero and two mismatches observed between primers and *ToBRFV*-strain genomes. Several binding sites were identified in genomes of *Tobamovirus* species closely-related to the *ToBRFV*; 15, 14, 11, and 9 primers produced partial matches to *Tobacco mosaic virus (TMV)*, *Tomato mottle mosaic virus (ToMMV)*, *Rehmannia mosaic virus* (*ReMV),* and *Tomato mosaic virus (ToMV)* genomes, respectively (Table 1). Fewer matches, between zero and six, were identified for the other nine *Tobamovirus* genomes which are more distantly related to the *ToBRFV* (Table 1; Salem et al., 2016).

**Table 1.** *In silico* Analysis of *ToBRFV*-specific Primer Binding to 192 *Tobamovirus* Genomes. *In silico* primer binding analysis using Benchling online software (*Benchling, 2023)* identified one potential binding site in 179 *ToBRFV* genomes for all 40 *ToBRFV*-specific primers. A limited number of *ToBRFV*-specific primers partially bound to other *Tobamovirus* genomes. A greater number of primer binding sites was observed in genomes of species sharing more recent ancestry with the *ToBRFV*.

| *Tobamovirus species name* | *Accession number* | *Primer Count* |
| --- | --- | --- |
| *All 179 ToBRFV genomes* | *Supplementary 1* | *40* |
| *Tobacco mosaic virus (TMV)* | *NC_001367.1* | *15* |
| *Tomato mottle mosaic virus (ToMMV)* | *NC_022230.1* | *9* |
| *Tomato mosaic virus (ToMV)* | *NC_002692.1* | *14* |
| *Rehmannia mosaic virus (ReMV)* | *NC_009642.1* | *11* |
| *Pepper mild mottle virus (PMMoV)* | *NC_003630.1* | *3* |
| *Bell pepper mottle tobamovirus (BPeMV)* | *NC_009642.1* | *6* |
| *Obuda pepper virus (ObPV)* | *NC_003852.1* | *1* |
| *Paprika mild mottle virus (PaMMV)* | *NC_004106.1* | *3* |
| *Tobacco middle green mosaic virus (TMGMV)* | *NC_001556.1* | *2* |
| *Cucumber green mottle mosaic virus (CGMMV)* | *NC_001801.1* | *0* |
| *Cucumber fruit mottle mosaic virus (CFMMV)* | *NC_002633.1* | *0* |
| *Brugmansia mild mottle virus (BrMMV)* | *NC_010944.1* | *6* |
| *Zucchini green mottle mosaic virus (ZGMMV)* | *NC_003878.1* | *0* |

Our *in silico* primer binding analysis suggests all 40 primers could amplify the genomic sequences of all 179 *ToBRFV*-strains listed on NextStrain while maintaining an overlapping binding scheme across each genome (Table 1, Figure 1). *In silico* results demonstrate that our primers are highly-specific to *ToBRFV* genome sequences yet remain non-specific at the strain-level. Moreover, limited-binding to other *Tobamovirus* genomes shows our primers are unlikely to amplify the complete genome of off-target species; however, partial genome segments could potentially be amplified.

### Read quality control and adaptor trimming

Sequenced RNA shotgun libraries produced on average 4.64 Gbp and 15 627 313 reads per sample, whereas *ToBRFV*-Seq libraries produced on average 179 820 Mbp and 597 554 reads per sample, in total 63.7 GB of data was produced (Table 2). Due to the low level of viral reads and high level of background eukaryotic and prokaryotic DNA found in environmental samples, shotgun prepared libraries often require a high sequencing depth to obtain even a modest amount of viral reads (Pérez-Cataluña et al., 2021; Pérez-Losada et al., 2020). Thus, RNA shotgun samples were loaded at a 15 × greater concentration than *ToBRFV*-Seq prepared samples. The majority of reads were high-quality, with RNA shotgun and *ToBRFV*-Seq prepared samples containing an average of 94.07% and 83.72% reads passing filters, respectively (Table 2).

**Table 2.** Total and Filtered Reads from RNA Shotgun and *ToBRFV*-Seq Prepared Sequencing Libraries. The total number of reads, number of filtered paired reads (QF Read Pairs), and percent of quality filtered (%PF) reads obtained from WWI samples and a nuclease-free water blank prepared with either RNA shotgun or *ToBRFV*-Seq. Samples A-E were averaged, excluding the NTC (no-template control) blank.

| Sample | RNA Shotgun |  |  |  | ToBRFV-Seq |  |  |  |
| --- | --- | --- | --- | --- | --- | --- | --- | --- |
|  | Total Gbp | Total Read Pairs | QF Read Pairs | % PF | Total Gbp | Total Read Pairs | QF Read Pairs | % PF |
| **Average** | **4.6400000** | **15 627 313** | **14 631 264** | **94.07** | **0.1798200** | **597 554** | **502 007** | **83.72** |
| A | 5.5000000 | 18 528 523 | 17 138 798 | 92.50 | 0.1688000 | 560 908 | 451 928 | 80.57 |
| B | 4.9000000 | 16 434 198 | 15 693 587 | 95.49 | 0.2103000 | 698 848 | 609 572 | 87.23 |
| C | 3.6000000 | 12 035 815 | 11 788 803 | 97.95 | 0.1589000 | 527 913 | 445 326 | 84.36 |
| D | 5.4000000 | 18 266 390 | 16 248 968 | 88.96 | 0.2054000 | 682 673 | 586 488 | 85.91 |
| E | 3.8000000 | 12 871 637 | 12 286 164 | 95.45 | 0.1557000 | 517 430 | 416 720 | 80.54 |
| Blank | 0.0396000 | 131 614 | 123 528 | 93.86 | 0.0000571 | 190 | 108 | 56.84 |

### Taxonomic profile of WWI samples prepared by RNA shotgun and *ToBRFV*-Seq methods

Domain-level taxonomic classification of read pairs by Kraken2 against the standard database was computed to compare the recovery of viral reads obtained with each library prep method (Figure 2-A,B; Wood et al., 2019). RNA shotgun reads were mostly bacterial, representing on average 89.40% of sample reads, 0.06% were viral, while the remaining 10.53% of reads were eukaryotic, archaeal, or other. Comparatively, on average, 97.38% of *ToBRFV*-Seq reads were classified as viral while the remaining 2.62% of reads were eukaryotic, viral, archaeal, or other (Figure 2-C). The high recovery of bacterial reads in cDNA shotgun libraries is likely a result of residual bacterial genomic DNA, rRNA, and mRNA sequences in prepared samples and is a common limitation of viral metagenomic sequencing (Elbehery & Deng, 2022; Guajardo-Leiva et al., 2020). DNA contamination could be prevented by including thorough quality checks to confirm DNA has been completely removed from the sample prior to library preparation steps (Guajardo-Leiva et al., 2020). Notably, DNA contamination did not impact the recovery of viral reads using our *ToBRFV*-Seq, even though the same extracts were used for each library prep method. Moreover, we have found DNase treatment unnecessary to obtain a high number of specific reads with our *ToBRFV*-Seq assay (data available upon request).

**Figure 2.**
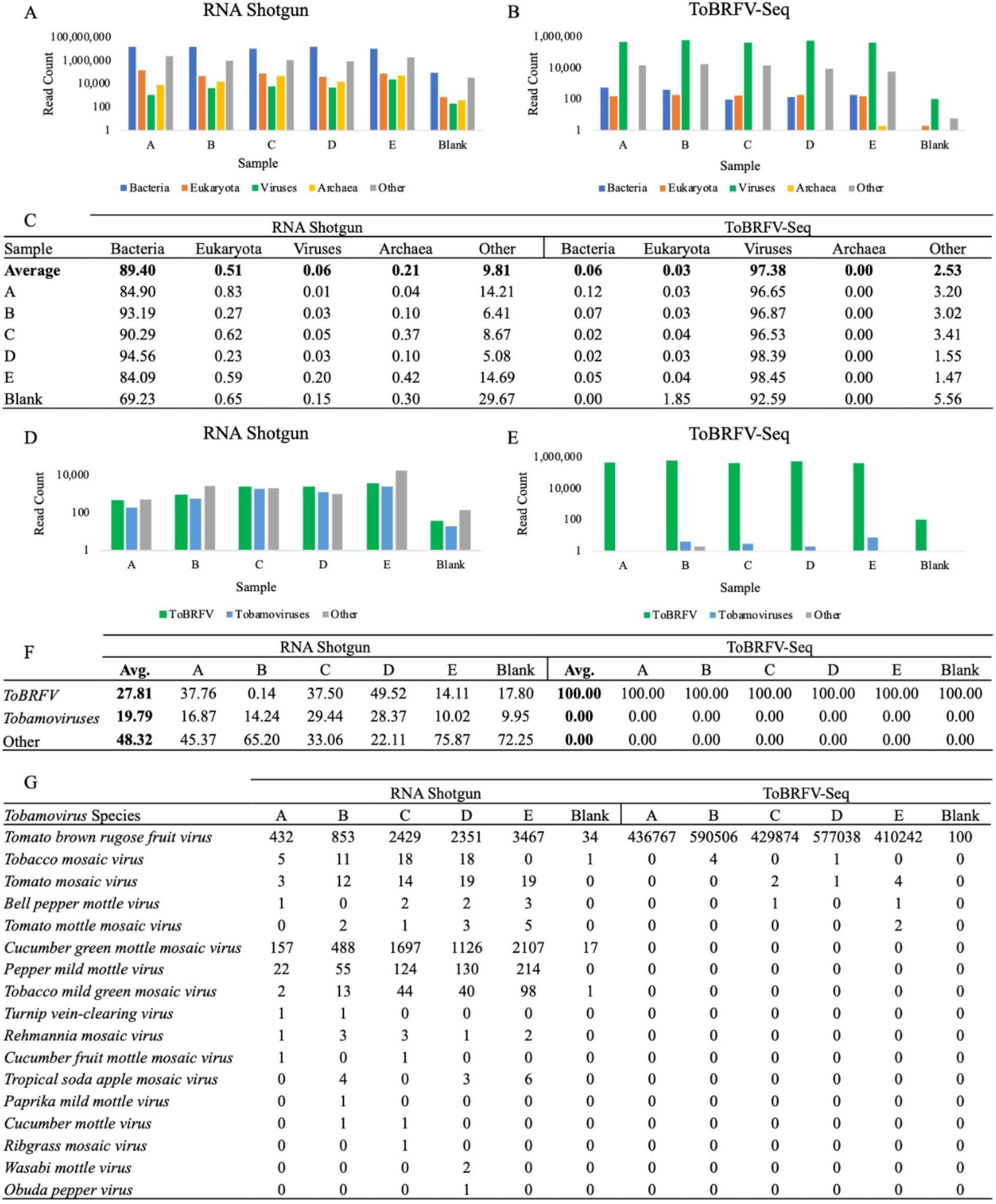
Taxonomic Classification of RNA Shotgun and *ToBRFV*-Seq Read Pairs at the Domain- and Species-level. Kraken2 classified read pair counts at the domain-level for (A) RNA shotgun library prepared samples, and (B) *ToBRFV*-Seq prepared samples (Wood et al., 2019). At the domain-level, categories included are viral, archaea, bacterial, eukaryotic, or other reads, where other includes unclassified reads, plasmids, adapters, linkers, and/or primers. (C) The percentage of reads represented by each domain for both prep methods. Samples A-E were used to calculate the average, the blank was excluded. (D&E) Percentage of Kraken2 classified *ToBRFV* (T), all other *Tobamoviruses* (OT), and the remainder of virus classified reads (RR) (F). Counts of *Tobamovirus* classified read pairs at the species-level (G). (A,B,D&E) Reads are represented on a log-scale to magnify and visualize low-abundance reads. Two-tailed tests were used to compare the mean read counts of RNA shotgun and *ToBRFV*-Seq classified reads, respectively. (D) RNA shotgun samples, *p* < 0.05, n = 5, df = 4, t-values, (T-OT) 3.569, (T-RR) -1.015, & (RR-OT) -1.181, critical t-value ±2.776. (E) *ToBRFV*-Seq samples, *p* < 0.05, n = 5, df = 4, t-values, (T-OT) 12.523, (T-RR) 12.523, & (RR-OT) 2.449, critical t-value ±2.776. Thus, there is a statistically significant difference between the number of *ToBRFV* and other *Tobamovirus* reads produced by RNA shotgun sequencing. Whereas, there is a statistically significant difference between the number of *ToBRFV* reads produced compared to both other Tobamovirus reads and the read remainder.

Viral read pairs were further classified at the species-level to evaluate the recovery of *ToBRFV*, other *Tobamovirus*, and all other virus species by each library preparation method (Figure 2-D&E). As expected, RNA shotgun sequencing yielded a mixture of *Tobamovirus* reads with a high proportion specific to the *ToBRFV*, as well as other virus species (Figure 2-D&F). On the other hand, our *ToBRFV*-Seq assay almost exclusively produced *ToBRFV* reads (Figure 2-E&F). Although other *Tobamoviruses* were observed in all WWI samples, they were not significantly enriched within *ToBRFV*-Seq samples (Figure 2-G). This demonstrates that our primers do not amplify these closely-related off-target sequences when present in low-abundance and are therefore highly-specific to *ToBRFV* genomes.

Of note, the *ToBRFV-*Seq blank sample, prepared using nuclease-free water, contained 100 *ToBRFV* read pairs (Figure 2-G). This is likely due to sample contamination prior to or during library preparation and is a commonly encountered issue when preparing tiled-amplicon sequencing assays (Mackie et al., 2022). Thus, extra precautions should be taken to reduce sample contamination during RNA extraction and PCR set up (Hirschhorn et al., 2023).

### Mapping reads to a *ToBRFV* reference genome

The quality of *ToBRFV* genome recovery was assessed by mapping reads from samples to a *ToBRFV* genome using bowtie2, and assessing coverage, completeness, and depth of read mapping to the reference. Our *ToBRFV*-specific primers target 95.7% of the *ToBRFV* genome sequence, forgoing 64 bp and 210 bp on the 5’ and 3’ genome ends, respectively. These regions are less than 400 bp and therefore too small to be included in our tiled amplicon primer scheme. Thus, the percentage of genome coverage was calculated for both the total and PCR-targeted genome regions.

Both library preparation methods were able to obtain nearly complete or complete coverage of the total and PCR-targeted genome, however, RNA shotgun samples contained far fewer reads and had a lower depth of coverage then *ToBRFV*-Seq samples (Table 3). Moreover, RNA shotgun samples on average contained 15 627 313 read pairs, yet, only 0.02 % of reads aligned to the *ToBRFV* reference genome (Table 2 & 3). Conversely, *ToBRFV*-Seq samples contained on average 597 554 read pairs with 99.86 % of reads aligning to the reference genome (Table 2 & 3). RNA shotgun samples utilized a much greater sequencing capacity then our *ToBRFV-*Seq samples, yet, still yielded fewer *ToBRFV* reads and lower depth of sequencing. Thus, our *ToBRFV*-Seq assay offers an alternative and more robust approach for sequencing *ToBRFV* genomes that is highly specific and requires minimal sequencing capacity. On average, *ToBRFV*-Seq samples covered 99.92 % of the PCR-targeted genome sequence with 0-6 bp missing on alignment terminal ends.

**Table 3.**
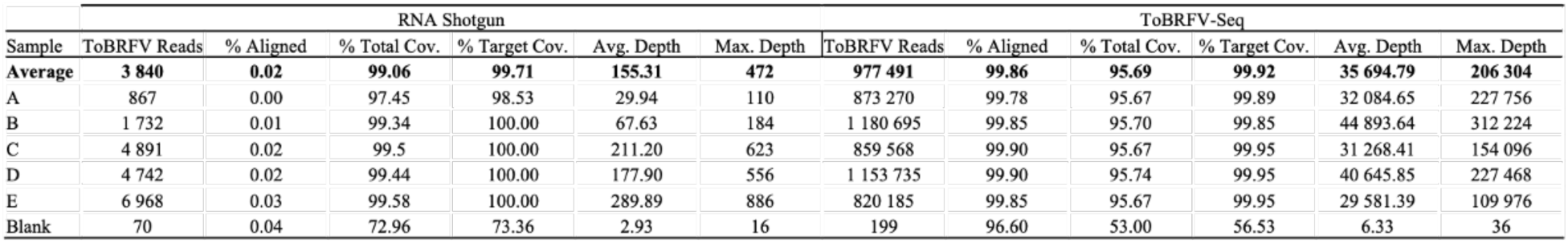
Alignment and Coverage of RNA Shotgun and *ToBRFV*-Seq Reads Mapped to a *ToBRFV* Genome. Number of *ToBRFV* reads and percentage of total reads that align to the *ToBRFV* genome NC_028478.1, % coverage of the total (% Total Cov) and PCR-targeted (% Target Cov) genomic region, the average read depth, and maximum read depth. Samples A-E were included in the average and the blank was excluded.

### Estimating the Relative Abundance of *ToBRFV* Clades in Wastewater Influent

We adapted our tool, Alcov, to estimate the abundance of *ToBRFV* clades in Illumina sequencing reads and named the adapted tool Altob. Clade-specific mutations were defined by constructing a Maximum Likelihood tree from unique *ToBRFV* isolates listed on the NextStrain 2022 build, clade-specific MSAs aligned to the reference genome NC_028478.1 were used to define mutations (Botermans et al., 2023). We were able to resolve seven of the eight clades identified on NextStrain, however, clade five isolates did not form a distinct clade and appeared in several divergent branches (Table S2, Figure S1). Benchmarking analysis of Altob was performed using synthetic Illumina read datasets of 1 000, 10 000, and 100 000 reads using genomes which represent each ToBRFV clade (Botermans et al., 2023).

Altob correctly identified all clades and estimated between 94.2% and 98.7% relative-abundance for the respective clades (Figure 3-A), except for clade five. Clade five contains the wildtype *ToBRFV* used to root our tree and define mutations in all other clades, therefore the clade five clade is not called by Altob (Botermans et al., 2023). Synthetic read datasets containing a mixture of clade-specific reads were also correctly identified using Altob (Figure 3-B). We expected a 50% abundance call for each clade in files containing two genomes, a 25% call in files containing four genomes, and a 12.5% call in files containing all eight genomes. Slight differences in the estimated and expected read abundance for some clades were observed (Figure 3-B), which can likely be attributed to the low-frequency of mutations in some strains and therefore absence of mutations in some randomly generated reads. Thus, Altob is well suited to identify and distinguish between *ToBRFV* clades with the exception of clade 5, however abundance scores should be treated as close estimates and not as exact measurements.

**Figure 3.**
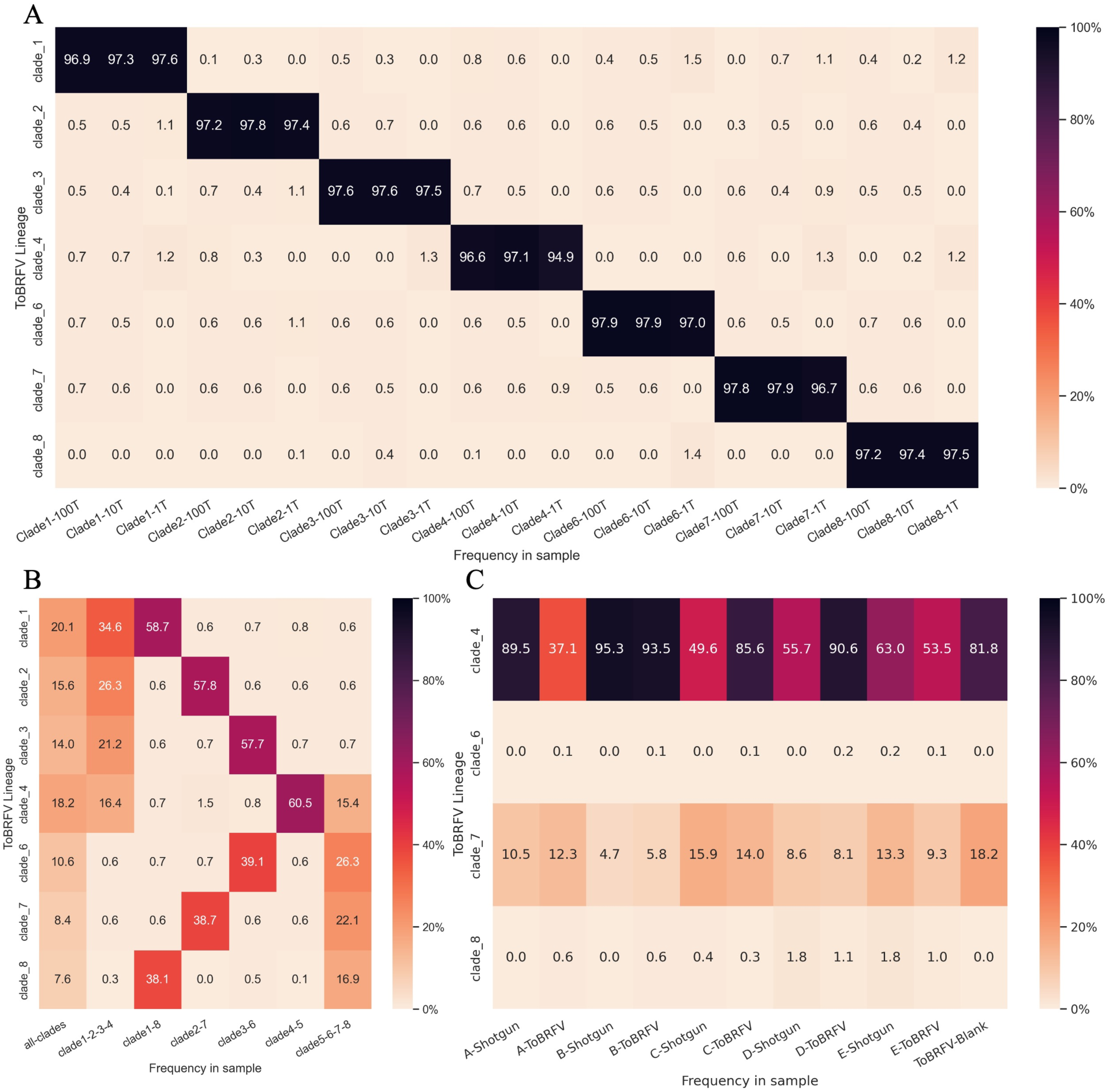
*ToBRFV* Clade-Abundance Estimates of Synthetic Read Datasets and WWI samples with Altob. Heatmaps display Altob clade-abundance estimates of (A) datasets containing 1 000, 10 000, or 100 000 synthetic reads from eight *ToBRFV* genomes representative of each clade. (B) datasets combining 100 000 synthetic reads from either all eight genomes, four genomes, or two genomes. (C) Reads from WWI samples prepared with RNA shotgun (Shotgun) or our *ToBRFV-*Seq assay (Tiled) (A-E).

Finally, Altob was used to evaluate the clades present in our WWI samples (Figure 3-C). In samples A-E, clades four and seven were identified in all RNA Shotgun and *ToBRFV-*Seq samples, demonstrating the wide-spread transmission of specific *ToBRFV*-clades in Ontario WWI. Clade four strains have been identified in North America, as such the identification of clade four strains in Ontario wastewater influent is unsurprising (Botermans et al., 2023). Clade seven strains have been identified in the Netherlands and Belgium, suggesting the transmission of these strains to Ontario either through the agricultural industry and/or the import of tomato products (Botermans et al., 2023; Rothman & Whiteson, 2022; Zhang et al., 2006). In each sample, the predicted clade abundance of clade 7 was relatively similar regardless of library preparation method with an average difference of ∓1.86% between RNA shotgun and *ToBRFV*-Seq estimates for each respective sample. However, clade 4 yielded different abundance calls when samples were prepared by each assay and had a average difference of ∓26.92%.

Generally, Altob is a robust prediction tool capable of estimating *ToBRFV* clade abundances in samples with as little as 1 000 *ToBRFV* reads. However, some differences in the estimated clades when using differing library preparation methods demonstrates that Altob predicted clade abundances should be treated as general estimates. The identification of additional *ToBRFV* strain genome sequences, particularly in clades 6 and 8 which each contain only three known strains, would enable more clade-specific mutations to be identified and added to Altob constellation files, thus improving Altob’s ability to classify samples by *ToBRFV* clade. Identifying and tracking strain-specific virus clades or lineages in WWI has proven a useful tool, particularly in SARS-CoV-2 pandemic preparedness efforts (Karthikeyan et al., 2022). Together, our *ToBRFV-*Seq assay and Altob, could be a valuable resource to identify clade-specific *ToBRFV* infections, explore potential transmission routes, and aid in combating transmission events.

### Assessment of viral shotgun reads at a species-level

To investigate the presence and abundance of viruses in WWI, viral reads from RNA shotgun samples were taxonomically classified at the species-level and the abundance of species assigned read-pairs was evaluated (Figure 4). In all five WWI samples, twenty-six viruses were consistently found which included plant infecting viruses, human pathogens, enteric bacteriophages, insect viruses, and amoeba infecting viruses (Table S4).

**Figure 4.**
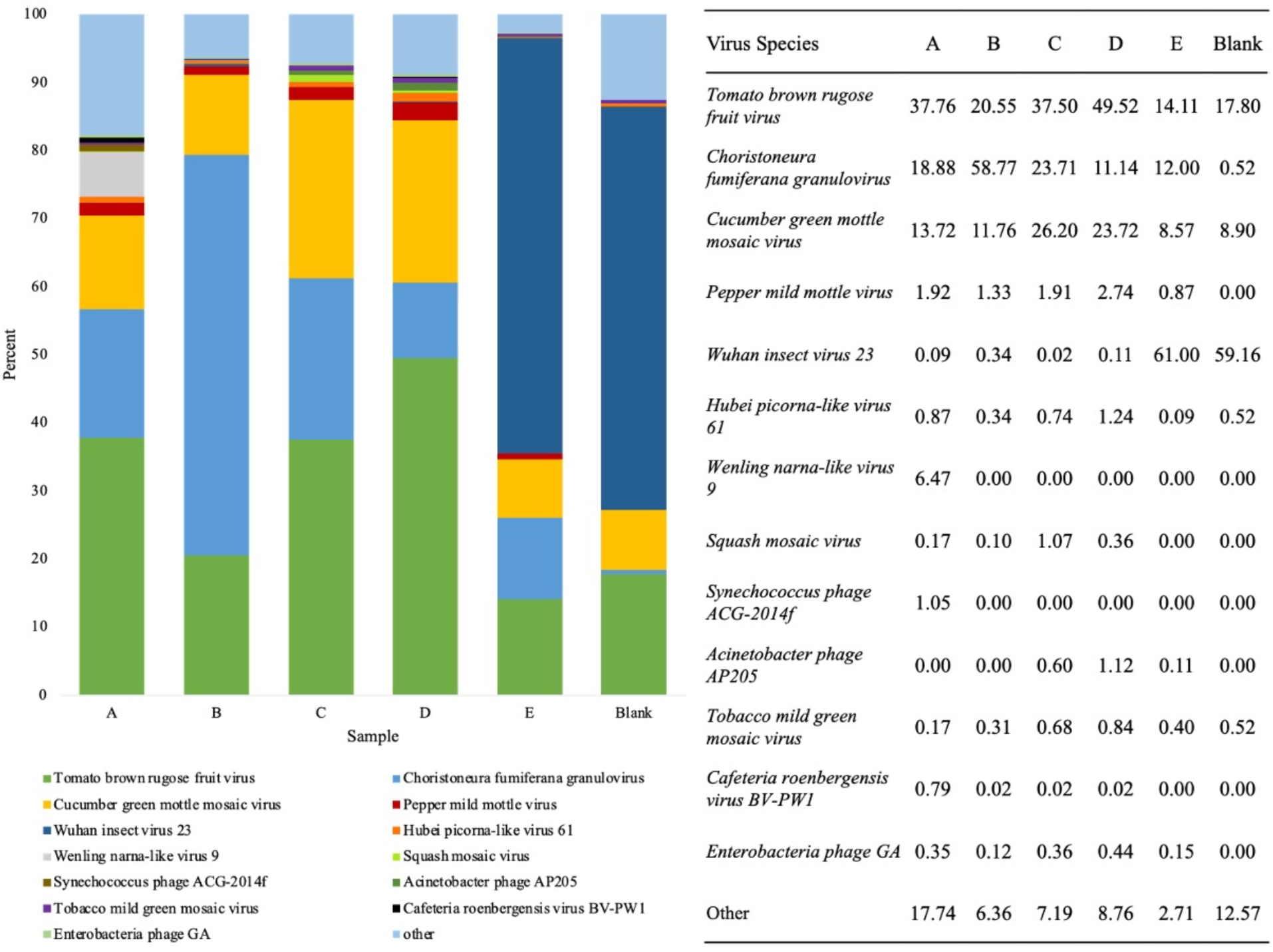
Evaluation of virus-species abundance in WWI samples using counts of viral reads taxonomically classified by Kraken2. (A) Percentage of virus paired-read counts of the seven most abundant virus-species found in each sample and count of all other species. (B) Percentage of virus-species classified reads assigned to the most abundant species in each WWI sample prepared by RNA shotgun and a blank.

The counts and percentage of paired-reads assigned to the seven most prevalent virus-species identified in each sample were compared to ascertain which species were consistently found in high-abundance (Figure 4). In samples A-D, either *ToBRFV* or *Choristoneura fumiferana granulovirus* (*ChfuGV)* read-pairs were in the highest abundance while *Cucumber green mottle mosaic virus (CGMMV)* and *Pepper mild mottle virus (PMMoV)* were on average the third and fourth most prevalent virus species, respectively (Figure 4-B). *ToBRFV, CGMMV,* and *PMMoV* are plant-pathogens belonging to the *Tobamovirus* genus, while *ChfuGV* is an insect-pathogen that infects budworms (Jones, 2021; Rashidan et al., 2008). In sample E, however, *Wuhan insect virus 23,* is the most-abundant single virus species detected. Preparation or sequencing biases may have resulted in the over-estimation of this virus in samples. However, these samples were collected during summer months, therefore there may have been a high insect to human virus transmission experienced by individuals in the population serviced by site E. Replicative data at this site could be used to discern between these and other potential possibilities.

The high relative abundance of *ToBRFV*-specific reads in RNA shotgun samples indicates that *ToBRFV* is widespread and highly prevalent in Ontario WWI (Figure 4). Moreover, comparing counts of *ToBRFV*-specific reads versus all other *Tobamovirus* reads with a one-tailed t-test (*p* < 0.05, n = 5, df = 4, t-values, t-value = 3.569, critical t-value ±2.776) demonstrates a statistically significant difference in read counts, with *ToBRFV* reads outnumbering all other *Tobamovirus* reads combined (Figure 2-D). The higher abundance of *ToBRFV* reads than *PMMoV* and all other *Tobamoviruses* is consistent with findings of Rothman & Whiteson, 2022, who investigated WWI from Californian treatment plants for the presence of eight *Tobamoviruses* using qPCR & Illumina sequencing. Of the eight *Tobamoviruses* viruses, their analysis identified the *ToBRFV* as the most abundant in California WWI, even more prevalent than both *PMMoV* and *CGMMV* (Rothman & Whiteson, 2022).

### Possible assay applications to determine ToBRFV transmission and prevalence

*PMMoV* is a commonly used human fecal indicator due to its reliable presence in human feces and has become a popular RT-qPCR standard to normalize cycle threshold (Ct) values from *SARS-CoV-2* assays (Zhang et al., 2006; Symonds et al., 2016; Torii et al., 2022; Karthikeyan et al., 2022). Since the *SARS-CoV-2* pandemic, a multitude of virus and chemical human fecal indicators, such as *PMMoV*, caffeine, and nicotine, have been recently evaluated (Ciannella et al., 2023; Maal-Bared et al., 2023; Oloye et al., 2023). However, a consensus on the best standard has not yet been reached and the reliability of *PMMoV* and other standards is quite variable between studies due to factors such as, population size, seasonal changes, diet, sample processing, and detection methods (Farkas et al., 2020; Hsu et al., 2022; Kitajima et al., 2018; Oloye et al., 2023; Rainey et al., 2023; Torii et al., 2022). Although *PMMoV* was found in all our RNA shotgun samples, it was consistently found in a lower abundance relative to *ToBRFV* suggesting *ToBRFV* may be more readily detectable, and thus be a more robust fecal indicator and RT-qPCR standard than *PMMoV* (Figure 4). Indeed, two studies examining human infant oropharyngeal and fecal viromes within the first year of life identified the *ToBRFV* by metagenomic sequencing in infant fecal (Aguado-García et al., 2020) and infant oropharyngeal (Rivera-Gutiérrez et al., 2023) viromes. Moreover, a recent comparative study of *PMMoV* and *ToBRFV* as fecal contamination indicators showed *ToBRFV* is more prevalent in wastewater and matched the established microbial tracking marker cross-assembly phage (crAssphage) abundance (Natarajan et al, 2023). This suggests that *ToBRFV* may be a suitable marker replacement to *PMMoV* in wastewater. *ToBRFV* has likely been overlooked as a fecal indicator because *PMMoV* was first introduced as a fecal indicator in 2006 long before the first recorded *ToBRFV* outbreak in 2014 (Luria et al., 2017; Salem et al., 2016; Zhang et al., 2006).

### Effectiveness of and possible improvements to viral capture and *ToBRFV*-Seq procedures

Microbiome A particles, manufactured by Ceres Nanosciences™, are magnetic capture beads which offer a fast and easy solution to isolate numerous virion species from wastewater, including, *SARS-CoV-2*, *Influenza*, *Respiratory syncytial virus*, *PMMoV*, *Norovirus*, and others (Ahmed et al., 2023; Karthikeyan et al., 2022; https://www.ceresnano.com/microbiome). We utilized Microbiome A particles to recover *ToBRFV* sequences from all of our WWI samples (Figure 2) and to our knowledge, this is the first demonstration of *ToBRFV* capture and concentration using Microbiome A particles.

Virus sequence yields in samples concentrated with Microbiome A particles provide a rapid, and user-friendly method to isolate *ToBRFV* and other viruses for targeted sequencing methods. However, Microbiome A beads may not be the best option for virus metagenomic RNA shotgun sequencing, due to the high contamination of bacterial DNA sequences. Thus, when performing viral metagenomic analysis a combination of ultracentrifugation and filtration or other robust viral isolation methods should be utilized (Ferandez-Cassi et al., 2018; Hjelmsø et al., 2017).

## Conclusions

To aid in *ToBRFV* detection and ongoing surveillance efforts, we designed and developed a *ToBRFV* genome tiled-amplicon sequencing assay that is robust, specific, and efficient. Moreover, we designed a computational tool, Altob, that estimates the relative-abundance of *ToBRFV*-clades in Illumina paired reads. Additionally, we demonstrate that Microbiome A particles are an effective method for isolating *ToBRFV* from WWI. Lastly, we demonstrate that *ToBRFV* is widespread and one of the most abundant virus species in Ontario, Canada wastewater influent. Due to its stability and abundance in WWI, the *ToBRFV* has the potential to be a useful indicator species in wastewater and/or used as a RT-qPCR standard to quantify viral content in wastewater, and more broadly, used in other wastewater epidemiology efforts and human fecal studies, however this would require further study (Natarajan et al, 2023). Moreover, due to the transmissibility of several *Tobamovirus* species in reclaimed wastewater, including *PMMoV* (Bačnik et al., 2020), the closely related *ToBRFV* may also be transmissible in reclaimed wastewater used in crop irrigation which could facilitate transmission of active *ToBRFV* virions to crops causing viral outbreaks.

## Data Availability and Links

NextStrain *ToBRFV* 2022 build access - https://nextstrain.nrcnvwa.nl/ToBRFV/20220412

Altob software access - https://github.com/Ellmen/altob

Raw read data with human reads removed available under BioProject PRJNA1030775.

## Acknowledgments

We are grateful to have received our wastewater samples through the Ontario Wastewater Surveillance Initiative led by the Ontario Ministry of Environment Conservation and Parks. We would like to thank the Public Health Units Peel Public Health, York Region Public Health, Region of Waterloo Public Health and Paramedic Services, and Kingston, Frontenac and Lennox & Addington Public Health as well as the members of Dr. Mark Servos’ research laboratory at the University of Waterloo and of Dr. Stephen Brown’s research laboratory for coordinating the delivery of the specific samples used in this research. We would like to thank the Natural Science and Engineering Research Council, the Ontario Ministry of Environment Conservation and Parks, and Mitacs for providing funding support for this project.

## References

Abrahamian, P., Cai, W., Nunziata, S. O., Ling, K., Jaiswal, N., Mavrodieva, V. A., Rivera, Y., & Nakhla, M. K. (2022). Comparative Analysis of Tomato Brown Rugose Fruit Virus Isolates Shows Limited Genetic Diversity. Viruses, 14(2816). 10.3390/v14122816

Adams, M. J., Adkins, S., Bragard, C., Gilmer, D., Li, D., MacFarlane, S. A., Wong, S.-M., Melcher, U., Ratti, C., Ryu, K. H., & ICTV Report Consortium. (2017). ICTV Virus Taxonomy Profile:Virgaviridae. Journal of General Virology, 98(8), 1999–2000. 10.1099/jgv.0.000884

Adams, M. J., Antoniw, J. F., & Kreuze, J. (2009). Virgaviridae: a new family of rod-shaped plant viruses. Archives of Virology, 154(12), 1967–1972. 10.1007/s00705-009-0506-6

Aguado-García, Y., Taboada, B., Morán, P., Rivera-Gutiérrez, X., Serrano-Vázquez, A., Iša, P., Rojas-Velázquez, L., Pérez-Juárez, H., López, S., Torres, J., Ximénez, C., & Arias, C. F. (2020). Tobamoviruses can be frequently present in the oropharynx and gut of infants during their first year of life. Scientific Reports, 10(13595). 10.1038/s41598-020-70684-w

Ahmed, W., Bivins, A., Korajkic, A., Metacalfe, S., Smith, W. J.M., & Simpson, S. L. (2023). Comparative analysis of Adsorption-Extraction (AE) and Nanotrap® Magnetic Virus Particles (NMVP) workflows for the recovery of endogenous enveloped and non-enveloped viruses in wastewater. Science of the Total Environment, 859. 10.1016/j.scitotenv.2022.160072

ARTIC Network. (2020). artic-ncov2019. artic-network. Retrieved May 7, 2023, from https://github.com/artic-network/artic-ncov2019

ARTIC Network. (2023, April 26). ARTIC Mulitplex PCR (Full pathogen list). ARTIC Network. Retrieved May 7, 2023, from https://community.artic.network/t/artic-multiplex-pcr-full-pathogens-list/494

Bačnik, K., Kutnjak, D., Pecman, A., Mehle, N., Žnidarič, M. T., Aguirre, I. G., & Ravnikar, M. (2020). Viromics and infectivity analysis reveal the release of infective plant viruses from wastewater into the environment. Water Research, 177(115628). 10.1016/j.watres.2020.115628

Bai, G., Lin, S., Hsu, Y., & Chen, S. (2022). The Human Virome: Viral Metagenomics, Relations with Human Diseases, and Therapeutic Applications. Viruses, 14(278). 10.3390/v14020278

Benchling [Biology Software]. (2023). https://benchling.com

Bolger, A. M., Lohse, M., & Usadel, B. (2014). Trimmomatic: A flexible trimmer for Illumina sequence data. Bioinformatics, 30(15), 2114–2120. 10.1093/bioinformatics/btu170

Botermans, M., de Koning, P. P.M., Oplaat, C., Fowkes, A. R., McGreig, S., Skelton, A., Adams, I. P., Fox, A., De Jonghe, K., Demers, J. E., Roenhorst, J. W., Westenberg, M., & van de Vossenberg, B. T.L.H. (2023). Tomato Brown Rugose Fruit Virus Nextstrain Build Version 3: Rise of a Novel Clade. PhytoFrontiers, 0(0), 0. 10.1094/phytofr-09-22-0090-a

Christou, A., Agüera, A., Bayona, J. M., Cytryn, E., Fotopoulos, V., Lambropoulou, D., Manaia, C. M., Michael, C., Revitt, M., Schröder, P., & Fatta-Kassions, D. (2017). The potential implications of reclaimed wastewater reuse for irrigation on the agricultural environment: The knowns and unknowns of the fate of antibiotics and antibiotic resistant bacteria and resistance genes – A review. Water Research, 123, 448–467. 10.1016/j.watres.2017.07.004

Ciannella, S., González-Fernández, C., & Gomez-Pastora, J. (2023). Recent progress on wastewater-based epidemiology for COVID-19 surveillance: A systematic review of analytical procedures and epidemiological modeling. Science of the Total Environment, 878. 10.1016/j.scitotenv.2023.162953

Danecek, P., Bonfield, J., Liddle, J., Marshall, J., Ohan, V., Pollard, M. O., Whitwham, A., Keane, T., McCarthy, S. A., Davies, R. M., & Li, H. (2021). Twelve years of SAMtools and BCFtools. GigaScience, 10, 1–4. 10.1093/gigascience/giab008

Edgar, R. C. (2004). MUSCLE: multiple sequence alignment with high accuracy and high throughput. Nucleic Acid Research, 32, 1792–1797. 10.1093/nar/gkh340

Ellmen, I., Lynch, M. D.J., Nash, D., Cheng, J., Nissimov, J. I., & Charles, T. C. (2021). Alcov: Estimating Variant of Concern Abundance from SARS-CoV-2 Wastewater Sequencing Data. medRxiv. doi10.1101/2021.06.03.21258306

Eren, A. M., Kiefl, E., Shaiber, A., Veseli, I., Miller, S. E., Schechter, M. S., Fink, I., Pan, J. N., Yousef, M., Fogarty, E. C., Trigodet, F., Watson, A. R., Esen, Ö. C., Moore, R. M., Clayssen, Q., Lee, M. D., Kivenson, V., Graham, E. D., Merrill, B. D., … Willis, A. D. (2021). Community-led, integrated, reproducible multi-omics with anvi’o. Nature Microbiology, 6, 3–6. 10.1038/s41564-020-00834-3

Farkas, K., Walker, D. I., Adriaenssens, E. M., McDonald, J. E., Hillary, L. S., Malham, S. K., & Jones, D. L. (2020). Viral indicators for tracking domestic wastewater contamination in the aquatic environment. Water Research, 181. 10.1016/j.watres.2020.115926

Ferandez-Cassi, X., Timoneda, N., Martínez-Puchol, S., Rusiñol, M., Rodriguez-Manzano, J., Figuerola, N., Bofill-Mas, S., Abril, J. F., & Girones, R. (2018). Metagenomics for the study of viruses in urban sewage as a tool for public health surveillance. Science of the Total Environment, 618, 870–880. 10.1016/j.scitotenv.2017.08.249

Fidan, H., Sarikaya, P., Yildiz, K., Topkaya, B., Erkis, G., & Calis, O. (2021). Robust molecular detection of the new Tomato brown rugose fruit virus in infected tomato and pepper plants from Turkey. Journal of Integrative Agriculture, 20(8), 2170–2179. 10.1016/S2095-3119(20)63335-4

Gibbs, A. (1999). Evolution and origins of tobamoviruses. The Royal Society, 354, 593–602.

Goh, P. S., Ahmad, N. A., Lim, J. W., Liang, Y. Y., Kang, H. S., Ismail, A. F., & Arthanareeswaran, G. (2022). Microalgae-Enabled Wastewater Remediation and Nutrient Recovery through Membrane Photobioreactors: Recent Achievements and Future Perspective. Membranes, 12(1094). 10.3390/membranes12111094

Gouy, M., Tannier, E., Comte, N., & Parsons, D. P. (2021). Seaview Version 5: A Multiplatform Software for Multiple Sequence Alignment, Molecular Phylogenetic Analyses, and Tree Reconciliation. Methods in Molecular Biology, 2231. 10.1007/978-1-0716-1036-7_15

Hak, H., & Spiegelman, Z. (2021). The Tomato Brown Rugose Fruit Virus Movement Protein Overcomes Tm-22 Resistance in Tomato While Attenuating Viral Transport. Molecular Plant-Microbe Interactions, 34(9), 1024–1032. 10.1094/MPMI-01-21-0023-R

Hill, S. C., Neto de Vasconcelos, J., Granja, B. G., Théze, J., Jandondo, D., Neto, Z., Mirandela, M., dos Santos Sebastião, C., Cândido, A. L.M., Clemente, C., Pereria da Silva, S., de Oliverira, T., Pybus, O. G., Faria, N. R., & Afonso, J. M. (2019). Early Genomic Detection of Cosmopolitan Genotype of Dengue Virus Serotype 2, Angola, 2018. Emerging infectious Diseases, 25(4). 10.3201/eid2504.180958

Hirschhorn, J. W., Avery, A., & Schandl, C. A. (2023). Managing a PCR Contamination Event Pathology Laboratory. Methods in Molecular Biology, 2621, 15–26. 10.1007/978-1-0716-2950-5_2

Hjelmsø, M. H., Hellmér, M., Fernandez-Cassi, X., Timoneda, N., Lukjancenko, O., Seidel, M., Elsässer, D., Aarestrup, F. M., Löfström, C., Bofill-Mas, S., Abril, J. F., Girones, R., & Schultz, A. C. (2017). Evaluation of Methods for the Concentration and Extraction of Viruses from Sewage in the Context of Metagenomic Sequencing. PLoS ONE, 12(1). 10.1371/journal.pone.0170199

Hsu, S.-Y., Bayati, M., Li, C., Hsieh, H.-Y., Belenchia, A., Klutts, J., Zemmer, S. A., Reynolds, M., Semkiw, E., Johnson, H.-Y., Foley, T., Wieberg, C. G., Wenzel, J., Johnson, M. C., & Lin, C.-H. (2022). Biomarkers selection for population normalization in SARS-CoV-2 wastewater-based epidemiology. Water Research, 223. 10.1016/j.watres.2022.118985

Jones, R. A.C. (2021). Global Plant Virus Disease Pandemics and Epidemics. Plants, 10(233). doi10.3390/plants10020233

Kareninen, L., Ogola, J., Kivistö, I., Smura, T., Aaltonen, K., Jääskeläinen, A. J., Kibiwot, S., Masika, M. M., Nyaga, P., Mwaengo, D., Anzala, O., Vapalahti, O., Webala, P. W., Forbes, K. M., & Sironen, T. (2020). Range Expansion of Bombali Virus in Mops condylurus Bats, Kenya, 2019. Emerging Infectious Disease, 26(12), 3007–3010. 10.3201/eid2612.202925

Karthikeyan, S., Levy, J. I., De Hoff, P., Humphrey, G., Birmingham, A., Jepsen, K., Farmer, S., Tubb, H. M., Valles, T., Tribelhorn, C. E., Tsai, R., Aigner, S., Sathe, S., Moshiri, N., Henson, B., Mark, A. M., Hakim, A., Baer, N. A., Barber, T., … Knight, R. (2022). Wastewater sequencing reveals early cryptic SARS-CoV-2 variant transmission. Nature, 609(7925), 101–108. 10.1038/s41586-022-05049-6

Kitajima, M., Sassi, H. P., & Torrey, J. R. (2018). Pepper mild mottle virus as a water quality indicator. ngj Clean Water, 1(19).

Kumar, A., Murthy, S., & Kapoor, A. (2017). Evolution of selective-sequencing approaches for virus discovery and virome analysis. Virus Research, 239, 172–179. 10.1016/j.virusres.2017.06.005

Langmead, B., & Salzberg, S. (2012). Fast gapped-read alignment with Bowtie 2. Nature Methods, 9(4), 357–359. 10.1038/nmeth.1923

Lefkowitz, E. J., Dempsey, D. M., Hendrickson, R. C., Orton, R. J., Siddell, S. G., & Smith, D. B. (2018). Virus taxonomy: the database of the International Committee on Taxonomy of Viruses (ICTV). Nucleic Acids Research, 46. 10.1093/nar/gkx932

Luria, N., Smith, E., Reingold, V., Bekelman, I., Lapidot, M., Levin, I., Elad, N., Tam, Y., Sela, N., Abu-Ras, A., Ezra, N., Haberman, A., Yitzhak, L., Lachman, O., & Dombrovsky, A. (2017). A New Israeli Tobamovirus Isolate InfectsTomato Plants Harboring Tm-22* Resistance Genes. PLOS ONE, 12(1). doi:10.1371/journal.pone.0170429

Ma, Z., Ding, M., Zhang, Z., Yang, X., & Zhou, X. (2021). Molecular characterization and pathogenicity of an infectious cDNA clone of tomato brown rugose fruit virus. Phytopathology Research, 3(14). 10.1186/s42483-021-00091-0

Maal-Bared, R., Qiu, Y., Li, Q., Gao, T., Hrudey, S. E., Bhava, S., Ruecker, N. J., Ellehoj, E., Lee, B. E., & Pang, X. (2023). Does normalization of SARS-CoV-2 concentrations by Pepper Mild Mottle Virus improve correlations and lead time between wastewater surveillance and clinical data in Alberta (Canada): comparing twelve SARS-CoV-2 normalization approaches. Science of the Total Environment, 856. 10.1016/j.scitotenv.2022.158964

Mackie, J., Kinoti, W. M., Chahal, S. I., Lovelock, D. A., Campbell, P. R., Tran-Nguyen, L. T.T., Rodoni, B. C., & Constable, F. E. (2022). Targeted Whole Genome Sequencing (TWG-Seq) of Cucumber Green Mottle Mosaic Virus Using Tiled Amplicon Multiplex PCR and Nanopore Sequencing. Plants, 11(2716), 10.3390/plants11202716.

Mehle, N., Bačnik, K., Bajde, I., Brodarič, J., Fox, A., Gutiérrez-Aguirre, I., Kitek, M., Kutnjak, D., Loh, Y. L., Ferreira, O. M.C., Ravnikar, M., Vogel, E., Vos, C., & Vučurović, A. (2023). Tomato brown rugose fruit virus in aqueous environments - survival and significance of water-mediated transmission. Frontiers in Plant Science, 14(1187920). 10.3389/fpls.2023.1187920

Milne, I., Stephen, G., Bayer, M., Cock, P. J. A., Pritchard, L., Cardle, L., Shaw, P. D., & Marshall, D. (2013). Using tablet for visual exploration of second-generation sequencing data. Briefings in Bioinformatics, 14(2), 193–202. 10.1093/bib/bbs012

Natarajan, A., Fremin, B. J., Schmidtke, D. T., Wolfe, M. K., Zlitni, S., Graham, K. E., Brooks, E. F., Severyn, C. J., Sakamoto, K. M., Lacayo, N. J., Kuersten, S., Koble, J., Caves, G., Kaplan, I., Singh, U., Jagannathan, P., Rezvani, A. R., Bhatt, A. S., & Boehm, A. B. (2023). Tomato brown rugose fruit virus Mo gene is a novel microbial source tracking marker. bioRxiv, 10.1101/2023.01.09.523366

Ochar, K., Ko, H., Woo, H., Hahn, B., & Hur, O. (2023). Pepper Mild Mottle Virus: An Infectious Pathogen in Pepper Production and a Potential Indicator of Domestic Water Quality. Viruses, 15(282). 10.3390/v15020282

Oloye, F. F., Xie, Y., Challis, J. K., Femi-Oloye, O. P., Brinkmann, M., McPhedran, K. N., Jones, P. D., Servos, M. R., & Giesy, J. P. (2023). Understanding common population markers for SARS-CoV-2 RNA normalization in wastewater – A review. Chemosphere, 333. 10.1016/j.chemosphere.2023.138682

Pérez-Cataluña, A., Cuevas-Ferrando, E., Randazzo, W., & Sánchez, G. (2021). Bias of library preparation for virome characterization in untreated and treated wastewaters. Science of the Total Environment, 767. 10.1016/j.scitotenv.2020.144589

Pérez-Losada, M., Arenas, M., Galán, J. C., Bracho, A., Hillung, J., García-González, N., & Conzález-Candelas, F. (2020). High-throughput sequencing (HTS) for the analysis of viral populations. Infection, Genetics and Evolution, 80(104208), 1567–1348. 10.1016/j.meegid.2020.104208

Quick, J., Grubaugh, N. D., Pullan, S. T., Claro, I. M., Smith, A. D., Gangavarapu, K., Oliveria, G., Robles-Sikisaka, R., Rogers, T. F., Beutler, N. A., Burton, D. R., Lewis-Ximenez, L. L., Goes de Jesus, J., Giovanetti, M., Hill, S. C., Black, A., Bedford, T., Carroll, M. W., Nunes, M., … Loman, N. J. (2017). Multiplex PCR method for MinION and Illumina sequencing of Zika and other virus genomes directly from clinical samples. Nature Protocols, 12(6), 1261–1276. 10.1038/nprot.2017.066

Rainey, A. L., Liang, S., Bisesi, J. H. J., Sabo-Attwood, T., & Maurelli, A. T. (2023). A multistate assessment of population normalization factors for wastewater-based epidemiology of COVID-19. PLOS ONE, 18(4).

Rashidan, K., Guertin, C., & Cabana, J. (2008). Granulovirus. Encyclopedia of Entomology. 10.1007/978-1-4020-6359-6_1155

Rencher, A. C., & Christensen, W. F. (2012). Methods of Multivariate Analysis (3rd ed.). Wiley.

Rivera-Gutiérrez, X., Morán, P., Taboada, B., Serrano-Vázquez, A., Isa, P., Rojas-Velázquez, L., Pérez-Juárez, H., López, S., Torres, J., Ximénez, C., & Arias, C. F. (2023). The fecal and oropharyngeal eukaryotic viromes of healthy infants during the first year of life are personal. Scientific Reports, 13(938). 10.1038/s41598-022-26707-9

Rizzo, D., Lio, D. D., Panattoni, A., Salemi, C., Cappellini, G., Bartolini, L., & Parrella, G. (2021). Rapid and Sensitive Detection of Tomato Brown Rugose Fruit Virus in Tomato and Pepper Seeds by Reverse Transcription Loop-Mediated Isothermal Amplification Assays (Real Time and Visual) and Comparison With RT-PCR End-Point and RT-qPCR Methods. Frontiers in Microbiology, 12(640932). 10.3389/fmicb.2021.640932

Rothman, J. A., & Whiteson, K. L. (2022). Sequencing and Variant Detection of Eight Abundant Plant-Infecting Tobamoviruses across Southern California Wastewater. Microbiology Spectrum. 10.1128/spectrum.03050-22

Salem, N., Mansour, A., Falk, B. W., & Turina, M. (2016). A new tobamovirus infecting tomato crops in Jordan. Archives in Virology, 161(2), 503–506. 10.1007/s00705-015-2677-7

Sandybayev, N., Beloussov, V., Strochkov, V., Solomadin, M., Granica, J., & Yegorov, S. (2022). Next Generation Sequencing Approaches to Characterize the Respiratory Tract Virome. Microorganisms, 10(2327). 10.3390/microorganisms10122327

Statistics Canada. (n.d.). Table 32-10-0456-01 Production and value of greenhouse fruits and vegetables. Statistics Canada. Retrieved May 25, 2023, from https://www150.statcan.gc.ca/t1/tbl1/en/cv.action?pid=3210045601

Symonds, E. M., Sinigalliano, C., Gidley, M., Ahmed, W., McQuig-Ulrich, S. M., & Breitbart, M. (2016). Faecal pollution along the southeastern coast of Florida and insight into the use of pepper mild mottle virus as an indicator. Applied Microbiology, 121(5), 1469–1481. 10.1111/jam.13252

Torii, S., Oishi, W., Zhu, Y., Thakali, O., Malla, B., Yu, Z., Zhao, B., Arakawa, C., Kitajima, M., Hata, A., Ihara, M., Kyuwa, S., Sano, D., Haramoto, E., & Katayama, H. (2022). Comparison of five polyethylene glycol precipitation procedures for the RT-qPCR based recovery of murine hepatitis virus, bacteriophage phi6, and pepper mild mottle virus as a surrogate for SARS-CoV-2 from wastewater. Science of the Total Environment, 807. 10.1016/j.scitotenv.2021.150722

van de Vossenberg, B., Visser, M., Bruinsma, M., Koenraadt, H., Westenberg, M., & Botermans, M. (2020). Real-time tracking of Tomato brown rugose fruit virus (ToBRFV) outbreaks in the Netherlands using Nextstrain. PLoS ONE, 15(10). 10.1371/journal.pone.0234671

van de Vossenburg, B., Dawood, T., Woźny, M., & Botermans, M. (2021). First Expansion of the Public Tomato Brown Rugose Fruit Virus (ToBRFV) Nextstrain Build; Inclusion of New Genomic and Epidemiological Data. PhytoFrontiers, 1(4), 359–363. 10.1094/phytofr-01-21-0005-a

Wood, D. E., Lu, J., & Langmead, B. (2019). Improved metagenomic analysis with Kraken 2. Genome Biology, 20(257). 10.1186/s13059-019-1891-0

Yang, Y., Walls, S. D., Gross, S. M., Schroth, G. P., Jarman, R. G., & Hang, J. (2018). Targeted sequencing of respiratory viruses in clinical specimens for pathogen identification and genome-wide analysis. Methods in Molecular Biology, 1838, 125–140. 10.1007/978-1-4939-8682-8_10

Zhang, S., Griffiths, J., Marchand, G., Bernards, M., & Wang, A. (2022). Tomato brown rugose fruit virus: An emerging and rapidly spreading plant RNA virus that threatens tomato production worldwide. Molecular Plant Pathology, 23(9), 1262–1277. 10.1111/mpp.13229

Zhang, T., Breitbart, M., Lee, W. H., Run, J., Wei, C. L., Soh, S. W.L., Hibberd, M. L., Liu, E. T., Rohwer, F., & Ruan, Y. (2006). RNA viral community in human feces: Prevalence of plant pathogenic viruses. PLoS Biology, 4(1), 0108–0118. 10.1371/journal.pbio.0040003

