## Supplemental tables and figures for "A Novel Tiled-Amplicon Sequencing Assay Targeting the *Tomato Brown Rugose Fruit Virus* (*ToBRFV*) Genome Reveals Widespread Distribution in Municipal Wastewater Treatment Systems in the Province of Ontario, Canada"

**Table S1.** List of sample sites, type of collection system, serviced population size, and raw data accessions.

| Sample | Region | Collection Site | Collection System | Population | Sample Type | Collection Date | Prep. Date | RNA-Seq data accession | *ToBRFV*-Seq data accession |
|---|---|---|---|---|---|---|---|---|---|
| A | Kingston | Cataraqui Bay | Wastewater Treatment Plant | 52 084 | 24-h composite | June 28th 2023 | July 17th 2023 | SAMN37915769 | SAMN37915775 |
| B | Peel | GE Booth | Wastewater Treatment Plant | 1 089 738 | 24-h composite | July 10th 2023 | July 17th 2023 | SAMN37915770 | SAMN37915776 |
| C | Cambridge | Cambridge | Wastewater Treatment Plant | 89 714 | 24-h composite | July 10th 2023 | July 17th 2023 | SAMN37915771 | SAMN37915777 |
| D | York | Warden - 407 | Linear Pipe Collection Site | 650 303 | Grab | July 10th 2023 | July 17th 2023 | SAMN37915772 | SAMN37915778 |
| E | York | Leslie Street | Pumping Station | 295 232 | Grab | July 10th 2023 | July 17th 2023 | SAMN37915773 | SAMN37915779 |

**Table S2.** Accession numbers of 179 ToBRFV Strains obtained from NCBI. *ToBRFV* genomes listed on NextStrain used to design primers and phylogenetic analysis. The strains used for tree building (PhyML Tree), assigned clades (Clade), and genomes used for read simulation (Sim. Reads) are listed. Unique strain genomes used in our PhyML tree are denoted yes (PhyML Tree), identical genomes are denoted no and the representative genome used in the PhyML Tree is listed (Rep. Genome). Strains were assigned to the same clade by both our PhyML tree and the NextStrain build, with the expectation of some clade 5 strains. Clade 5 strains predicted by NextStrain but not our analysis are labeled 5(N).

| Accession | PhyML Tree | Rep. Genome | Clade | Sim. Reads |
|---|---|---|---|---|
| OM515230 | yes | - | 3 | - |
| OM515231 | yes | - | - | - |
| OM515232 | yes | - | - | - |
| OM515233 | yes | - | 8 | - |
| OM515234 | yes | - | - | - |
| OM515235 | yes | - | 8 | - |
| OM515236 | yes | - | 3 | yes |
| OM515237 | yes | - | - | - |
| OM515238 | yes | - | 3 | - |
| OM515239 | yes | - | 1 | yes |
| OM515240 | yes | - | 3 | - |
| OM515241 | yes | - | 1 | - |
| OM515242 | yes | - | 3 | - |
| OM515243 | no | MW314135 | 7 | yes |
| OM515244 | yes | - | 3 | - |
| OM515245 | yes | - | 7 | - |
| OM515246 | yes | - | 7 | - |
| OM515247 | yes | - | 7 | - |
| OM515248 | no | OM515255 | 7 | - |
| OM515249 | no | OM515252 | 7 | - |
| OM515250 | yes | - | - | - |
| OM515251 | no | OM515252 | 7 | - |
| OM515252 | yes | - | 7 | - |
| OM515253 | no | OM515255 | 7 | - |
| OM515254 | yes | - | 7 | - |
| OM515255 | yes | - | 7 | - |
| OM515256 | yes | - | - | - |
| OM515257 | no | OM515266 | - | - |
| OM515258 | yes | - | 8 | yes |
| OM515259 | yes | - | 7 | - |
| OM515260 | no | OM515245 | 7 | - |
| OM515261 | yes | - | 1 | - |
| OM515262 | yes | - | - | - |
| OM515263 | yes | - | 3 | - |
| OM515264 | yes | - | 6 | - |
| OM515265 | yes | - | 7 | - |
| OM515266 | yes | - | - | - |
| OM515267 | yes | - | 3 | - |
| OM515268 | no | OM515255 | 7 | - |
| OM515269 | no | OM515245 | 7 | - |
| OM515270 | yes | - | 1 | - |
| OM515271 | yes | - | 7 | - |
| OM515272 | yes | - | 7 | - |
| OM718702 | yes | - | 7 | - |
| OM718703 | yes | - | 7 | - |
| OM718704 | yes | - | 7 | - |
| OM718705 | no | OM515255 | 7 | - |
| OM718706 | yes | - | 1 | - |
| MZ945419 | no | MZ945420 | 1 | - |
| MZ945420 | yes | - | 1 | - |
| MW349655 | yes | - | 4 | - |
| MZ323110 | yes | - | 5 | - |
| MZ438228 | yes | - | 5 | - |
| MZ202349 | yes | - | 5 (N) | - |
| MZ004925 | yes | - | 5 (N) | - |
| MT018320 | yes | - | 5 (N) | - |
| MW314091 | yes | - | 5 | - |
| MW314092 | yes | - | 2 | yes |
| MW314093 | no | MN882033 | 2 | - |
| MW314094 | yes | - | 2 | - |
| MW314095 | no | MW314097 | 3 | - |
| MW314096 | no | MW314097 | 3 | - |
| MW314097 | yes | - | 3 | - |
| MW314098 | no | MW314097 | 3 | - |
| MW314099 | no | MW314101 | 3 | - |
| MW314100 | no | MW314101 | 3 | - |
| MW314101 | yes | - | 3 | - |
| MW314102 | no | MN882034 | 3 | - |
| MW314103 | yes | - | 3 | - |
| MW314104 | no | MW314105 | 5 | - |
| MW314105 | yes | - | 5 | - |
| MW314106 | yes | - | 1 | - |
| MW314107 | no | MW314109 | 5 | - |
| MW314108 | yes | - | 5 | - |
| MW314109 | yes | - | 5 | - |
| MW314110 | yes | - | 5 | - |
| MW314111 | yes | - | 8 | - |
| MW314112 | yes | - | 1 | - |
| MW314113 | yes | - | - | - |
| MW314114 | yes | - | 5 | - |
| MW314115 | no | MW314116 | - | - |
| MW314116 | yes | - | - | - |
| MW314117 | yes | - | - | - |
| MW314118 | yes | - | 1 | - |
| MW314119 | no | MW314120 | 1 | - |
| MW314120 | yes | - | 1 | - |
| MW314121 | yes | - | 3 | - |
| MW314122 | yes | - | 3 | - |
| MW314123 | yes | - | 6 | yes |
| KT383474 | yes | - | 5 | - |
| MW314124 | no | MW314125 | 1 | - |
| MW314125 | yes | - | 1 | - |
| MW314126 | yes | - | 3 | - |
| MW314127 | yes | - | 3 | - |
| MW314128 | yes | - | 1 | - |
| MW314129 | no | MW314136 | 3 | - |
| MW314130 | yes | - | 1 | - |
| MW314131 | yes | - | 1 | - |
| MW314132 | no | MW314137 | 1 | - |
| MW314133 | no | MW314135 | 7 | - |
| MW314134 | no | MW314135 | 7 | - |
| MW314135 | yes | - | 7 | - |
| MW314136 | yes | - | 3 | - |
| MW314137 | yes | - | 1 | - |
| MP872414 | no | KT383474 | 5 | - |
| MP872415 | no | KX619418 | 5 (N) | - |
| MP875816 | no | KT383474 | 5 | - |
| MP875817 | no | KX619418 | 5 (N) | - |
| MN549394 | yes | - | 4 | yes |
| MN549395 | yes | - | 4 | - |
| MN549396 | yes | - | 4 | - |
| MT002973 | yes | - | 4 | - |
| MT107885 | yes | - | 5 (N) | - |
| MT118666 | no | KX619418 | 5 (N) | - |
| MN882011 | yes | - | 3 | - |
| MN882012 | yes | - | 1 | - |
| MN882013 | yes | - | 1 | - |
| MN882014 | yes | - | 1 | - |
| MN882015 | yes | - | 1 | - |
| MN882016 | yes | - | 1 | - |
| MN882017 | yes | - | 1 | - |
| MN882018 | yes | - | 1 | - |
| MN882019 | no | MN882020 | 3 | - |
| MN882020 | yes | - | 3 | - |
| MN882021 | yes | - | 3 | - |
| MN882022 | no | MN882034 | 3 | - |
| MN882023 | no | MN882034 | 3 | - |
| MN882024 | no | MN882034 | 3 | - |
| MN882025 | no | MN882064 | 3 | - |
| MN882026 | no | MN882027 | 1 | - |
| MN882027 | yes | - | 1 | - |
| MN882028 | no | MN882029 | 3 | - |
| MN882029 | yes | - | 3 | - |
| MN882030 | no | MN882033 | 2 | - |
| MN882031 | no | MN882033 | 2 | - |
| MN882032 | no | MN882033 | 2 | - |
| MN882033 | yes | - | 2 | - |
| MN882034 | yes | - | 3 | - |
| MN882035 | no | MN882034 | 3 | - |
| MN882036 | yes | - | 3 | - |
| MN882037 | no | MN882040 | 1 | - |
| MN882038 | no | MN882040 | 1 | - |
| MN882039 | no | MN882040 | 1 | - |
| MN882040 | yes | - | 1 | - |
| MN882041 | yes | - | 1 | - |
| MN882042 | no | MN882043 | 6 | - |
| MN882043 | yes | - | 6 | - |
| MN882044 | no | MN882045 | 1 | - |
| MN882045 | yes | - | 1 | - |
| MN882046 | no | MN882047 | 1 | - |
| MN882047 | yes | - | 1 | - |
| MN882048 | yes | - | 3 | - |
| MN882049 | no | MN882050 | 3 | - |
| MN882050 | yes | - | 3 | - |
| MN882051 | yes | - | 1 | - |
| MN882052 | yes | - | 1 | - |
| MN882053 | yes | - | 1 | - |
| MN882054 | yes | - | 1 | - |
| MN882055 | yes | - | 1 | - |
| MN882056 | no | MN882015 | 1 | - |
| MN882057 | yes | - | 1 | - |
| MN882058 | yes | - | 1 | - |
| MN882059 | no | MN882060 | 1 | - |
| MN882060 | yes | - | 1 | - |
| MN882061 | no | MN882062 | 1 | - |
| MN882062 | yes | - | 1 | - |
| MN882063 | no | MN882064 | 3 | - |
| MN882064 | yes | - | 3 | - |
| MN815773 | yes | - | - | - |
| MN013187 | yes | - | 5 (N) | - |
| MN013188 | yes | - | 5 (N) | - |
| MN182533 | yes | - | 1 | - |
| MK648157 | yes | - | 5 (N) | - |
| MN167466 | yes | - | 3 (5) | - |
| MK319944 | yes | - | 4 | - |
| MK165457 | yes | - | 5 (N) | yes |
| MK133093 | yes | - | 5 (N) | - |
| MK133095 | yes | - | 3 | - |
| KX619418 | yes | - | 5 (N) | - |

**Table S3.** List of *ToBRFV*-Seq Primer Sequences and Concentration of Pooled Primers.

| Primer Name | Pool | Sequence | Size | %GC | Tm | [Pool] |
|---|---|---|---|---|---|---|
| ToBRFV_1_LEFT | 1 | TTTACAACTACAATGGCATACACACA | 26 | 34.62 | 59.95 | 0.5 μM |
| ToBRFV_1_RIGHT | 1 | ATTGGGCATACAGCAGTGAACA | 22 | 45.45 | 60.74 | 0.5 μM |
| ToBRFV_2_LEFT | 2 | AATGATGCAGATCCCGTACGGA | 22 | 50 | 61.79 | 0.5 μM |
| ToBRFV_2_RIGHT | 2 | CGGCGTAACAAACATGGACATT | 22 | 45.45 | 60.27 | 0.5 μM |
| ToBRFV_3_LEFT | 1 | TGCTATTGCATTGCACAGTATATACG | 26 | 38.46 | 60.56 | 0.5 μM |
| ToBRFV_3_RIGHT | 1 | TGCGCTGTAAAATTGCTCACTAT | 23 | 39.13 | 59.56 | 0.5 μM |
| ToBRFV_4_LEFT | 2 | ACACCTGGTTTTGTAAGTTTTCTAGG | 26 | 38.46 | 60.12 | 0.5 μM |
| ToBRFV_4_RIGHT | 2 | CTGGCAGTCACTCCGTTGATAA | 22 | 50 | 60.53 | 0.5 μM |
| ToBRFV_5_LEFT | 1 | GCACTTATCAAGCCAAGGCACT | 22 | 50 | 61.64 | 0.5 μM |
| ToBRFV_5_RIGHT | 1 | ATCAAGCACTGGCATATCCACC | 22 | 50 | 60.93 | 0.5 μM |
| ToBRFV_6_LEFT | 2 | CGCATTAGAAATCAGGGTGCCT | 22 | 50 | 61.19 | 0.5 μM |
| ToBRFV_6_RIGHT | 2 | CCTCTTGCCATTGAACCCTTCA | 22 | 50 | 61 | 0.5 μM |
| ToBRFV_7_LEFT | 1 | TGTCGCATTGGCACTTAAAGATT | 23 | 39.13 | 59.75 | 0.5 μM |
| ToBRFV_7_RIGHT | 1 | TGGTCGCAACATCTAAGACTCC | 22 | 50 | 60.27 | 0.5 μM |
| ToBRFV_8_LEFT | 2 | CGGTGTCGAACCTAGTCAAGAT | 22 | 50 | 60.02 | 0.5 μM |
| ToBRFV_8_RIGHT | 2 | CGGTACTAAGATTAGATCTTCCTCAAAAT | 29 | 34.48 | 59.77 | 0.5 μM |
| ToBRFV_9_LEFT | 1 | GCAGTGCTAAAGTCGTCCTAGT | 22 | 50 | 60.27 | 0.5 μM |
| ToBRFV_9_RIGHT | 1 | GAGTTTCCACCTCATCAACCTCT | 23 | 47.83 | 60.25 | 0.5 μM |
| ToBRFV_10_LEFT | 2 | TGGAGACACACAACAAATTCCATAC | 25 | 40 | 60.14 | 0.5 μM |
| ToBRFV_10_RIGHT | 2 | GCGATGATAGATACAGGTGTCGG | 23 | 52.17 | 60.92 | 0.5 μM |
| ToBRFV_11_LEFT | 1 | ACGGACGTCCATACAGTACATG | 22 | 50 | 60.08 | 0.5 μM |
| ToBRFV_11_RIGHT | 1 | CAGCATCATAGTTATTTAACATGGTGC | 27 | 37.04 | 59.89 | 0.5 μM |
| ToBRFV_12_LEFT | 2 | GATGCAGGGACCCAATAGCAAT | 22 | 50 | 60.94 | 0.5 μM |
| ToBRFV_12_RIGHT | 2 | TCTACTACCAAAGATGCAGTATTCTCA | 27 | 37.04 | 60.15 | 0.5 μM |
| ToBRFV_13_LEFT | 1 | TTGGAAAATTTGGTGGCGATGA | 22 | 40.91 | 59.55 | 0.5 μM |
| ToBRFV_13_RIGHT | 1 | TTTGCCTTGTGAGCTCACTGAA | 22 | 45.45 | 60.87 | 0.5 μM |
| ToBRFV_14_LEFT | 2 | TTCAAAGCGAATATCCGGCCTT | 22 | 45.45 | 60.86 | 0.5 μM |
| ToBRFV_14_RIGHT | 2 | TGAATGTTGTAACGTCCCCACT | 22 | 45.45 | 60.08 | 0.5 μM |
| ToBRFV_15_LEFT | 1 | ACAAGGCCACAGGAAAACTACT | 22 | 45.45 | 60.01 | 0.5 μM |
| ToBRFV_15_RIGHT | 1 | TGAACTCTTCTAAGTGATCCCAATCC | 26 | 42.31 | 60.8 | 0.5 μM |
| ToBRFV_16_LEFT | 2 | ACGTGATACATCATGACAGAGGG | 23 | 47.83 | 60.18 | 0.5 μM |
| ToBRFV_16_RIGHT | 2 | TGAACCAATATCTTATCAACCTTGGAGA | 28 | 35.71 | 60.88 | 0.5 μM |
| ToBRFV_17_LEFT | 1 | TGAGTTCATAGACTTGTCAAAATCAGAA | 28 | 32.14 | 59.87 | 0.5 μM |
| ToBRFV_17_RIGHT | 1 | CGCAGCAATTTTAACATTTCTAATATTAACGAG | 33 | 30.3 | 62 | 0.5 μM |
| ToBRFV_18_LEFT | 2 | GGTTTCAGTTCAAAGTCGTTCCA | 23 | 43.48 | 59.87 | 0.5 μM |
| ToBRFV_18_RIGHT | 2 | TCCTTCTCAACCCTAACTTTTAAACC | 26 | 38.46 | 59.67 | 0.5 μM |
| ToBRFV_19_LEFT | 1 | ACCGGGAAAAAGTTTAGTAGTAAAAGTG | 28 | 35.71 | 60.76 | 0.5 μM |
| ToBRFV_19_RIGHT | 1 | AGGATCTAGTACCGCATTGTACCT | 24 | 45.83 | 61.02 | 0.5 μM |
| ToBRFV_20_LEFT | 2 | GGCCGACCCTATAGAATTAATAAATTTATGT | 31 | 32.26 | 60.84 | 0.5 μM |
| ToBRFV_20_RIGHT | 2 | GGTGCAGAGGACCATTGTAAAC | 22 | 50 | 59.69 | 0.5 μM |

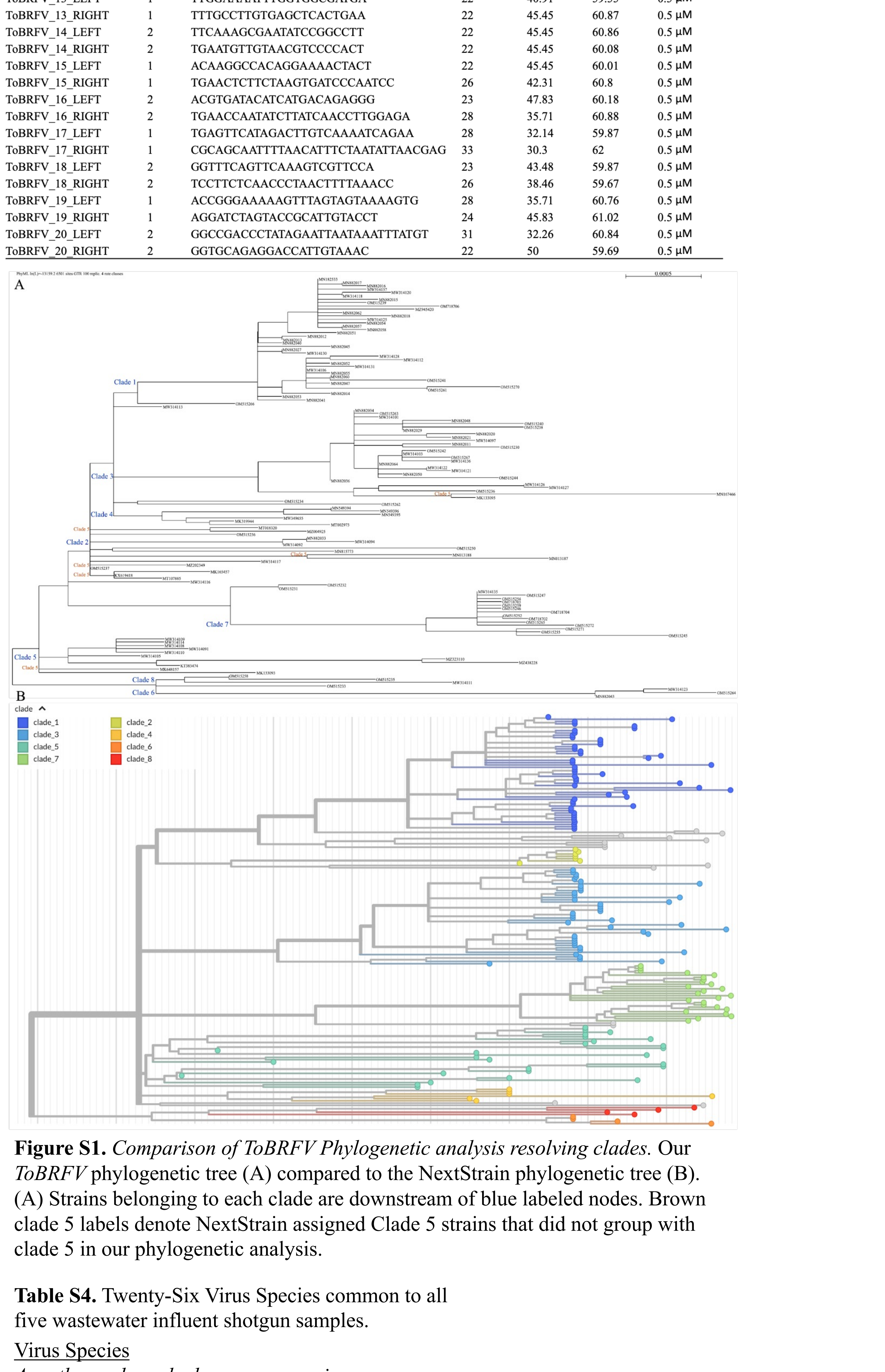


**Figure S1.** *Comparison of ToBRFV Phylogenetic analysis resolving clades.* Our *ToBRFV* phylogenetic tree (A) compared to the NextStrain phylogenetic tree (B). (A) Strains belonging to each clade are downstream of blue labeled nodes. Brown clade 5 labels denote NextStrain assigned Clade 5 strains that did not group with clade 5 in our phylogenetic analysis.

**Table S4.** Twenty-Six Virus Species common to all five wastewater influent shotgun samples.

| Virus Species |
|---|
| *Acanthamoeba polyphaga moumouvirus* |
| *Cactus virus X* |
| *Choristoneura fumiferana granulovirus* |
| *Cucumber green mottle mosaic virus* |
| *Emesvirus japonicum* |
| *Emesvirus zinderi* |
| *Enterobacteria phage GA* |
| *Enterobacteria phage Hgal1* |
| *Escherichia phage MS2* |
| *Garlic common latent virus* |
| *Garlic virus A* |
| *Hagavirus psychrophilum* |
| *Hubei picorna-like virus 61* |
| *Moumouvirus* |
| *Norwalk virus* |
| *Pepino mosaic virus* |
| *Pepper mild mottle virus* |
| *Pitaya virus X* |
| *Rehmannia mosaic virus* |
| *Schlumbergera virus X* |
| *Squash mosaic virus* |
| *Tobacco mild green mosaic virus* |
| *Tobacco mosaic virus* |
| *Tomato brown rugose fruit virus* |
| *Tomato mosaic virus* |
| *Wuhan insect virus 23* |
